# The *Ighmbp2*-R604X mouse recapitulates the severe SMARD1 clinical symptoms of aspiration, respiratory and feeding deficits

**DOI:** 10.1101/2025.08.27.672633

**Authors:** F. Javier Llorente Torres, Roxanne Muchow, Michelle Woolridge, Dennis Perez-Lopez, Catherine L. Smith, Nicole L. Nichols, Christian L. Lorson, Monique A. Lorson

## Abstract

Spinal muscular atrophy with respiratory distress type 1 (SMARD1) and Charcot Marie Tooth type 2S (CMT2S) are due to mutations in immunoglobulin mu binding protein two (*IGHMBP2*). We generated the *Ighmbp2*-R604X mouse (R605X-humans) to understand how alterations in IGHMBP2 function impact disease pathology. The IGHMBP2-R605X mutation is associated with patients with SMARD1 or CMT2S. The impact of this mutation is substantial, *Ighmbp2*^R604X/R604X^ mice have a decreased lifespan (6 days) and weight, and failure to thrive consistent with SMARD1 symptoms. Significant respiratory changes were present along with disease pathology of the phrenic nerve and diaphragm muscle fibers. *Ighmbp2*^R604X/R604X^ mice also presented with signs of milk aspiration and lung pathology. Interestingly, *Ighmbp2*^R604X/R604X^ mice were born with visible milk spots, but demonstrated reduction of the milk spot by P3, indicating deficits in suckling. Alterations in hindlimb electrophysiology were consistent with the sciatic nerve, hindlimb neuromuscular junction and muscle pathology. Injection of the ssAAV9-WT-*IGHMBP2* vector extended *Ighmbp2*^R604X/R604X^ survival a few days, due to reduced expression of the vector before death ensued. *Ighmbp2*^R604X/R604X^ phenotypes are consistent with the most severe SMARD1 clinical symptoms and for the first time a *Ighmbp2* mouse model demonstrates that milk aspiration and loss of the ability to suckle impact survival.

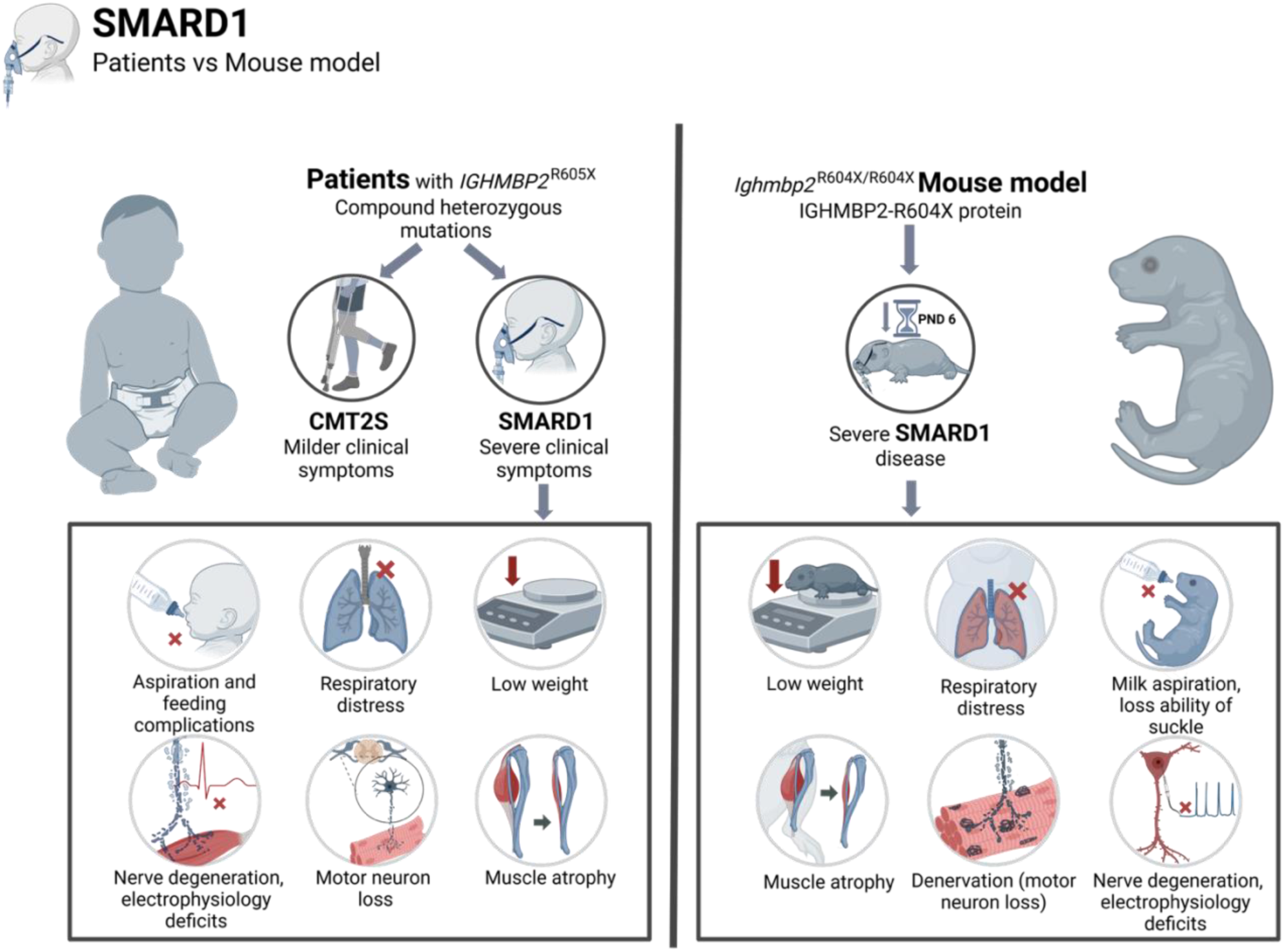

## Introduction

Mutations in immunoglobulin mu DNA binding protein two (*IGHMBP2*) give rise to two diseases: spinal muscular atrophy with respiratory distress (SMARD1) and Charcot Marie Tooth (CMT2S). SMARD1 is an autosomal recessive motor neuron disease that affects children during infancy, resulting in early death, and is distinguished from 5q-linked spinal muscular atrophy (SMA) by the genetic mutation and the clinical pathology. The primary clinical symptom, respiratory failure, is due to diaphragmatic paralysis, typically manifesting between 6 weeks to 13 months of age (1–6). SMARD1 clinical symptoms include distal lower limb muscle atrophy followed by proximal muscle weakness that results from anterior horn cell deterioration, intrauterine growth retardation, difficulties sucking, failure to thrive, autonomic nervous system and sensory defects, scoliosis and kyphosis (1–3, 5–7). SMARD1 presents with a variable disease onset, severity and symptoms. There is no clear patient correlation between the type of mutation and clinical presentation.

CMT2S patients present with a slow but progressive distal muscle weakness that manifests much later than SMARD1 and there is no respiratory impairment. Initial CMT2S symptoms include delayed milestones and difficulty walking. Nerve conduction studies show a reduction to absence of CMAP and sensory action potential, while deep tendon reflexes are nearly absent. There is typically no reduction in lifespan and patients must rely on palliative care (8–11).

IGHMBP2 is a member of the superfamily one (SF-1) DNA/RNA helicases that includes Up-frame shift 1(UPF1) and Senataxin (3, 12–17). Mutations in any of these helicases results in neurodegeneration despite having distinct cellular functions. While the function of IGHMBP2 that results in SMARD1/CMT2S has not been clearly defined, IGHMBP2 has proposed roles in immunoglobulin-class switching, pre-mRNA maturation, transcription regulation, pre-rRNA processing and translation (16, 18–21). More recent studies provide evidence for IGHMBP2 function within translation (22–24).

The *IGHMBP2* gene is comprised of 15 exons encoding 993 amino acids. Patient mutations are distributed throughout all 15 coding exons and within each of the proposed functional domains (helicase, R3H, zinc finger). Pathogenic variants in *IGHMBP2* occur in patients either as homozygous recessive or compound heterozygous mutations. We demonstrated that *IGHMBP2*-D565N and *IGHMBP2-*H924Y mutations result in diminished IGHMBP2 biochemical activity that correlates with disease severity and pathology (24, 25). Consistent with these studies, we demonstrated that the IGHMBP2-D565N mutation significantly reduces the binding affinity with activator of basal transcription (ABT1) while the IGHMBP2-H924Y mutation slightly alters IGHMBP2-ABT1 binding affinity (24).

The IGHMBP2-R605X mutation located in domain 2A of the IGHMBP2 helicase domain introduces a premature stop codon predicted to produce a truncated IGHMBP2 protein and loss of the IGHMBP2 R3H and zinc finger domains. Based on the Leiden open variation database (LOVD^3^), there are over 17 pathogenic truncating mutations within IGHMBP2; fourteen of those mutations are within the helicase domain (amino acids 19-641). We were interested in the IGHMBP2-R605X mutation as there are two reported patients with compound IGHMBP2-R605X mutations that result in SMARD1/respiratory distress at 91 and 365 days of life as well as three patients with compound IGHMBP2-R605X mutations with CMT2S clinical symptoms with symptoms initiating at 3+ years of life (4, 8, 26). LOVD^3^ also identified a single *IGHMBP2*^R605X/R605X^ patient that at six months that had respiratory failure and muscular hypotonia (27). To determine how the IGHMBP2-R605X mutation could generate such clinically diverse symptoms, we generated the orthologous mutation (R604X) in mice. In this manuscript, we report that the *Ighmbp*2-R604X (humans *IGHMBP2*-R605X) mice presented with some of the most severe symptoms observed in SMARD1 patients including failure to thrive, reduced suckling, aspiration of milk, severe respiratory distress and reduced motor function. Intracerebral ventricular (icv) injection of ssAAV9-*IGHMBP2* extended survival by a few days, likely due to the extent of disease pathology already present in *Ighmbp2*^R604X/R604X^ mice and the delayed expression of the ssAAV9-*IGHMBP2* vector.

## Results

### *Ighmbp2*^R604X/R604X^ mice had significantly reduced lifespan and weight with failure to thrive

IGHMBP2 is an RNA/DNA helicase of the SF1 family of helicases with a helicase core domain and C-terminal R3H and AN-1 zing finger (ZnF) domains (Figure 1A). The IGHMBP2-R605X mutation is located within the helicase domain (Figure 1A). We generated the orthologous *IGHMBP2*-R605X mutation (*Ighmbp2*-R604X) in FVB mice using CRISPR-Cas9, changing the nucleotide sequence at amino acid 604 from CGG to TAG (Figure 1B). To understand how the *Ighmbp2*-R604X mutation impacted disease, we bred *Ighmbp2*^+/R604X^ mice and examined survival and weight in *Ighmbp2*^+/R604X^, *Ighmbp2*^R604X/R604X^ and age-matched wild type mice. Survival was significantly reduced in *Ighmbp2*^R604X/R604X^ mice with an average lifespan of six days (*P*<0.0001); survival was not affected in *Ighmbp2*^+/R604X^ and wild type mice followed to weaning (Figure 1C-D). *Ighmbp2*^R604X/R604X^ mice did not demonstrate a significant difference in weight at birth (mean +/+=1.33 grams, +/R604X=1.37 grams, R604X/R604X=1.34 grams); however, there was a rapid decline in weight from P0 to P3 (mean +/+=2.56 grams, +/R604X=2.66 grams, R604X/R604X=1.76 grams) (Figure 1C, E). There was no statistical difference between age-matched wild type and *Ighmbp2*^+/R604X^ mice in weight. The mean weight at P6 was significantly different between wild type (4.35 grams ±0.66) and *Ighmbp2*^+/R604X^ (4.51 grams ±0.52) mice when compared to *Ighmbp2*^R604X/R604X^ (1.94 grams ±0.29) mice, but not between wild type and *Ighmbp2*^+/R604X^ mice (Figure 1C, E). These results show that *Ighmbp2*^R604X/R604X^ mice have a reduced lifespan and failure to thrive.

**Figure 1:**
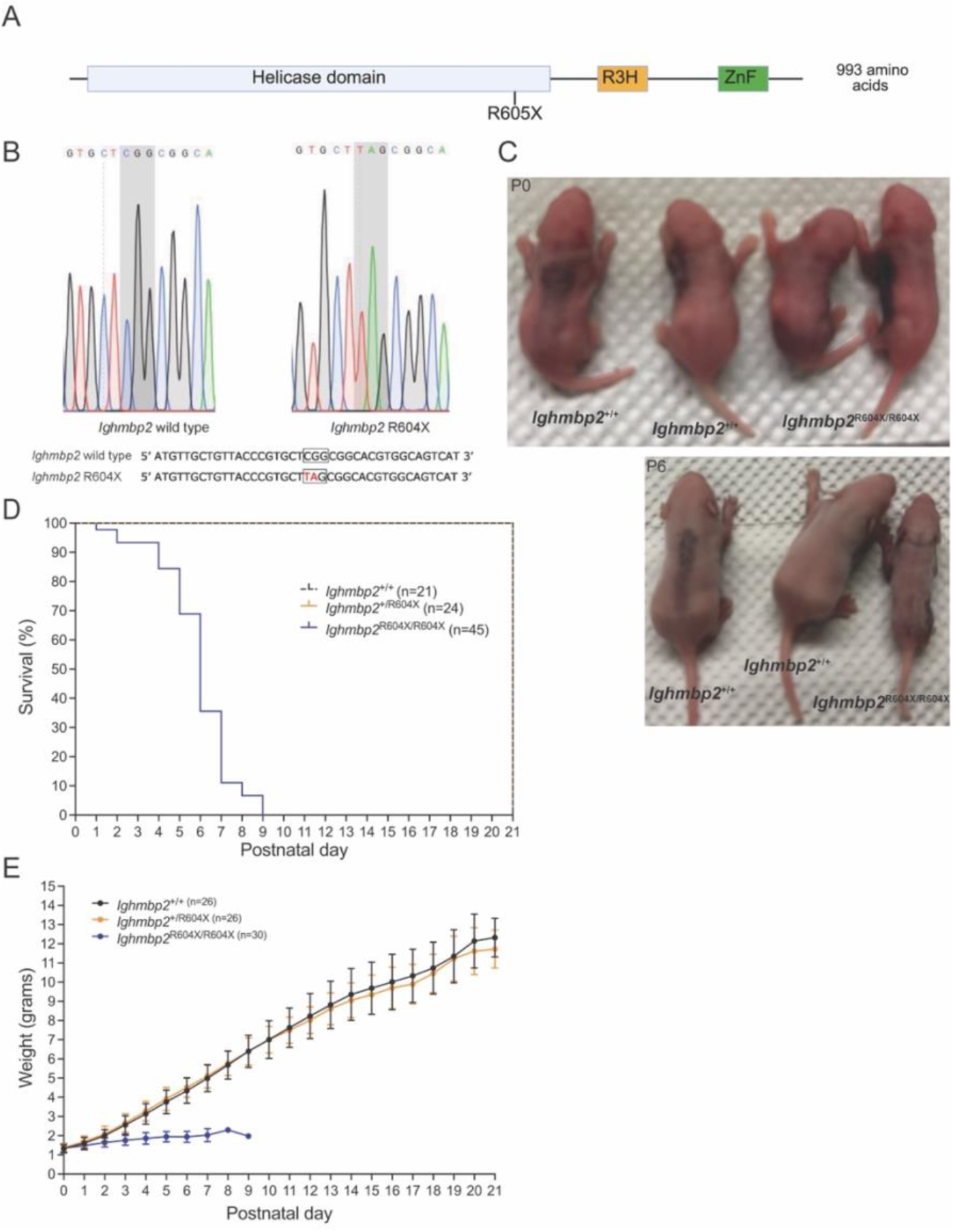
*Ighmbp2*^R604X/R604X^ mice had significantly reduced lifespan and weight. Wild type (black), *Ighmbp2*^+/R604X^ (orange), and *Ighmbp2*^R604X/R604X^ (blue). (**A**) A schematic representing the IGHMBP2 protein with the IGHMBP2-R605X mutation indicated (IGHMBP2-R604X in mice). (**B**) A schematic demonstrating the CGG to TAG genomic alteration of the mouse *Ighmbp2* gene to generate the *Ighmbp2*^+/R604X^ mouse. (**C**) Representative images of P0 (postnatal day) and P6 wild type, *Ighmbp2*^+/R604X^, and *Ighmbp2*^R604X/R604X^ mouse pups. (**D**) Survival curve for wild type, *Ighmbp2*^+/R604X^, and *Ighmbp2*^R604X/R604X^ mice followed for twenty-one days. Survival was twenty-one days for wild type and *Ighmbp2*^+/R604X^ mice while the mean survival for *Ighmbp2*^R604X/R604X^ mice was six days (*P*<0.0001). (**E**) Weight measured for twenty-one days. The mean weight was wild type=6.98 grams, *Ighmbp2*^+/R604X^=6.86 grams, *Ighmbp2*^R604X/R604X^=1.83 grams (*P*=0.0003 +/+ and R604X/R604X and *P*=0.0003 +/R604X and R604X/R604X). Statistical analyses: Survival data summary for survival, one-way ANOVA with Tukey’s multiple comparison for weight. Values are expressed as mean, n=number of mice.

### *Ighmbp2*^R604X/R604X^ mice showed rapid decline of milk spot

Due to the progressive loss of weight, we scored the presence of the milk sac/spot, an indicator of successful suckling by the pup. In the same animals, we compared survival, weight and the milk spot from P0 to P8 (Figure 2). Mice pups were given a score of 2.0 for a full milk spot, 1.0 for a partially full milk spot and 0 for no milk spot observed. By P3, there was a statistical difference between wild type (score 2.0) and *Ighmbp2*^R604X/R604X^ mice (score 1.2, *P*<0.0001) and *Ighmbp2*^+/R604X^ (score 1.9) and *Ighmbp2*^R604X/R604X^ mice (Figure 2C). At P4, when there was a significant weight difference (mean +/+=3.34 grams, +/R604X=3.01 grams, R604X/R604X=2.21 grams, *P*<0.0001), the milk spot score for *Ighmbp2*^R604X/R604X^ mice was statistically different (+/+=2.0, +/R604X=2.0, R604X/R604X=0.58, *P*<0.0001) (Figure 2B-C). Interestingly, all *Ighmbp2*^R604X/R604X^ mice analyzed had a visible milk spot at birth; however, the milk spot declined in most instances quite rapidly. There was a correlation between decreased survival, failure to gain weight and reduction of a milk spot (Figure 2A-D). These results suggest that the rapid reduction in lifespan and weight was at least in part due to the reduction/absence of nutrition associated with reduced suckling (reduction in milk spot).

**Figure 2:**
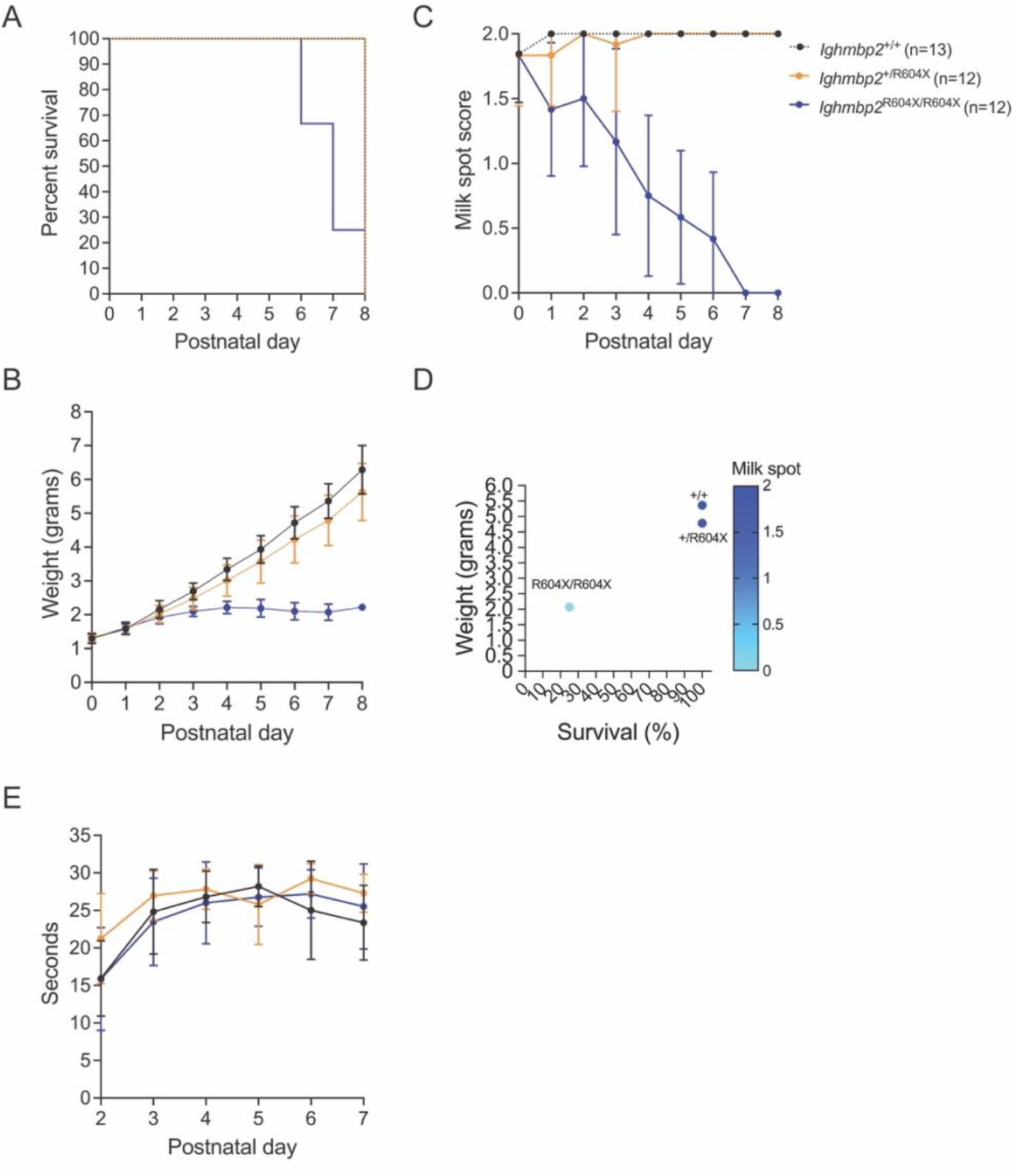
*Ighmbp2^R604X/R604X^* mice had a rapid decline of milk spot while maintaining latency to fall. Wild type (black), *Ighmbp2*^+/R604X^ (orange), and *Ighmbp2*^R604X/R604X^ (blue). A study of survival, weight, milk spot and latency to fall of same cohort of wild type, *Ighmbp2*^+/R604X^ and *Ighmbp2*^R604X/R604X^ mice for 8 days. (**A**) Survival curve with 25% survival at P7 for *Ighmbp2*^R604X/R604X^ mice. (**B**) Weight measured for eight days. P4 mean weight wild type=3.34 grams, *Ighmbp2*^+/R604X^=3.01 grams, *Ighmbp2*^R604X/R604X^=2.21 grams (*P*<0.0001 +/+ and R604X/R604X). (**C**) Presence of milk spot. Score of 2=full milk spot, 1=partial milk spot, 0=no milk spot present. P4 wild type=2, *Ighmbp2*^+/R604X^=2, *Ighmbp2*^R604X/R604X^=0.58 (*P*<0.0001 +/+ and R604X/R604X and *P*<0.0001 +/R604X and R604X/R604X). (**D**) Diagram showing correlation between survival, weight and presence of milk spot. (**E**) Latency to fall measured from P2-P7. Mean wild type=24.0 seconds, *Ighmbp2*^+/R604X^=26.4 seconds, *Ighmbp2*^R604X/R604X^=24.1 seconds (NS). Statistical analyses: Survival data summary for survival, one-way ANOVA with Tukey’s multiple comparison for weight and latency to fall, two-way ANOVA with Tukey’s multiple comparison for milk spot. Values are expressed as mean, n=number of mice, NS=not significant.

### Latency to fall in *Ighmbp2*^R604X/R604X^ mice was not affected

Since lifespan was dramatically reduced in *Ighmbp2*^R604X/R604X^ mice, we were unable to accomplish most motor function assessments; however, latency to fall was measured from P2-P7 (Figure 2E). There was no statistical difference in latency to fall between age-matched wild type, *Ighmbp2*^+/R604X^, and *Ighmbp2*^R604X/R604X^ mice (mean +/+=24.0 seconds, +/R604X=26.4 seconds, R604X/R604X=24.1 seconds); however, it was apparent that wild type and *Ighmbp2*^+/R604X^ had reduced latency to fall time due to their extensive movement as they reached P7. In contrast, *Ighmbp2*^R604X/R604X^ mice just hung from the tube and the reduced latency to fall was likely attributed to their reduced muscle strength (Figure 2E).

### *Ighmbp2*^R604X/R604X^ mice presented with extensive changes in respiration and lung pathology

SMARD1 patients present with respiratory changes early in life. The SMARD1 *Ighmbp2*^D564N/D564N^ mouse showed significant respiratory changes and did not respond to altered CO_2_ and O_2_ conditions (28). Interestingly, the short-lived *Ighmbp2*^D564N/H922Y^ cohort developed respiratory differences later and mounted a response to altered CO_2_ and O_2_ conditions (25). To determine whether respiration significantly impacted survival in *Ighmbp2*^R604X/R604X^ mice, we performed head-out plethysmography at P2 using customized pup chambers due to the reduced size and weight of the animals. Respiration was analyzed under three conditions normoxia (21% O_2_ + 0% CO_2_ + 79% N_2_), hypercapnia (7% CO_2_, 21% O_2_, balanced N_2_) (HC), and hypercapnia (7% CO_2_) + hypoxia (10.5% O_2_, balanced N_2_) (HC+HX). The following eight parameters were measured: frequency (f), tidal volume (VT), minute ventilation (VE), mean inspiratory flow (VT/TI), peak inspiratory flow (PIF), peak expiratory flow (PEF), inspiratory time (Ti) and expiratory time (Te) (Figure 3). *Ighmbp2*^R604X/R604X^ mice experienced significant respiratory changes when compared to wild type or *Ighmbp2*^+/R604X^ mice under all parameters measured except inspiratory time (Figure 3). There were no significant respiratory differences between wild type and *Ighmbp2*^+/R604X^ mice under any conditions (Figure 3). Respiratory frequency was significantly decreased in *Ighmbp2*^R604X/R604X^ mice when compared to wild type or *Ighmbp2*^+/R604X^ mice under all conditions (Figure 3A). Tidal volume was similar under normoxia conditions for all mice measured; however, unlike wild type and *Ighmbp2*^+/R604X^ mice, *Ighmbp2*^R604X/R604X^ mice did not respond to elevated CO_2_ (hypercapnia) nor reduced O_2_ and elevated CO_2_ conditions (HC+HX) (Figure 3B). Minute ventilation, which accounts for frequency and tidal volume, affirms that *Ighmbp2*^R604X/R604X^ mice experience significant respiratory changes and unlike wild type and *Ighmbp2*^+/R604X^ mice do not mount a response to altered CO_2_ and O_2_ conditions (Figure 3C). These results suggest that chemoreception is altered in *Ighmbp2*^R604X/R604X^ mice. To understand if respiration was impacted by respiratory muscle strength or respiratory obstruction we measured flow rates. Mean inspiratory flow (MIF-average flow rate of air during inspiration) was similar between all mice under normoxia; however, MIF was significantly decreased in *Ighmbp2*^R604X/R604X^ mice under hypercapnia and hypercapnia + hypoxia conditions (Figure 3D). Peak inspiratory flow, the maximum airflow achieved during inspiration, was significantly different in *Ighmbp2*^R604X/R604X^ mice under all three conditions (N, HC, HC+HX) affirming that *Ighmbp2*^R604X/R604X^ mice showed difficulty in the inspiratory phase of respiration (Figure 3E). When peak expiratory flow (PEF) was measured for wild type and *Ighmbp2*^+/R604X^ mice there were no significant differences under any conditions; however, *Ighmbp2*^R604X/R604X^ mice showed significantly reduced PEF under hypercapnia and hypercapnia + hypoxia suggesting *Ighmbp2*^R604X/R604X^ mice also showed difficulty in the expiratory phase of respiration (Figure 3F). *Ighmbp2*^R604X/R604X^ mice did not show differences in inspiratory time under any conditions but there was significantly prolonged expiratory time under normoxia that was reduced under challenged respiratory conditions (Figure 3G-H).

**Figure 3:**
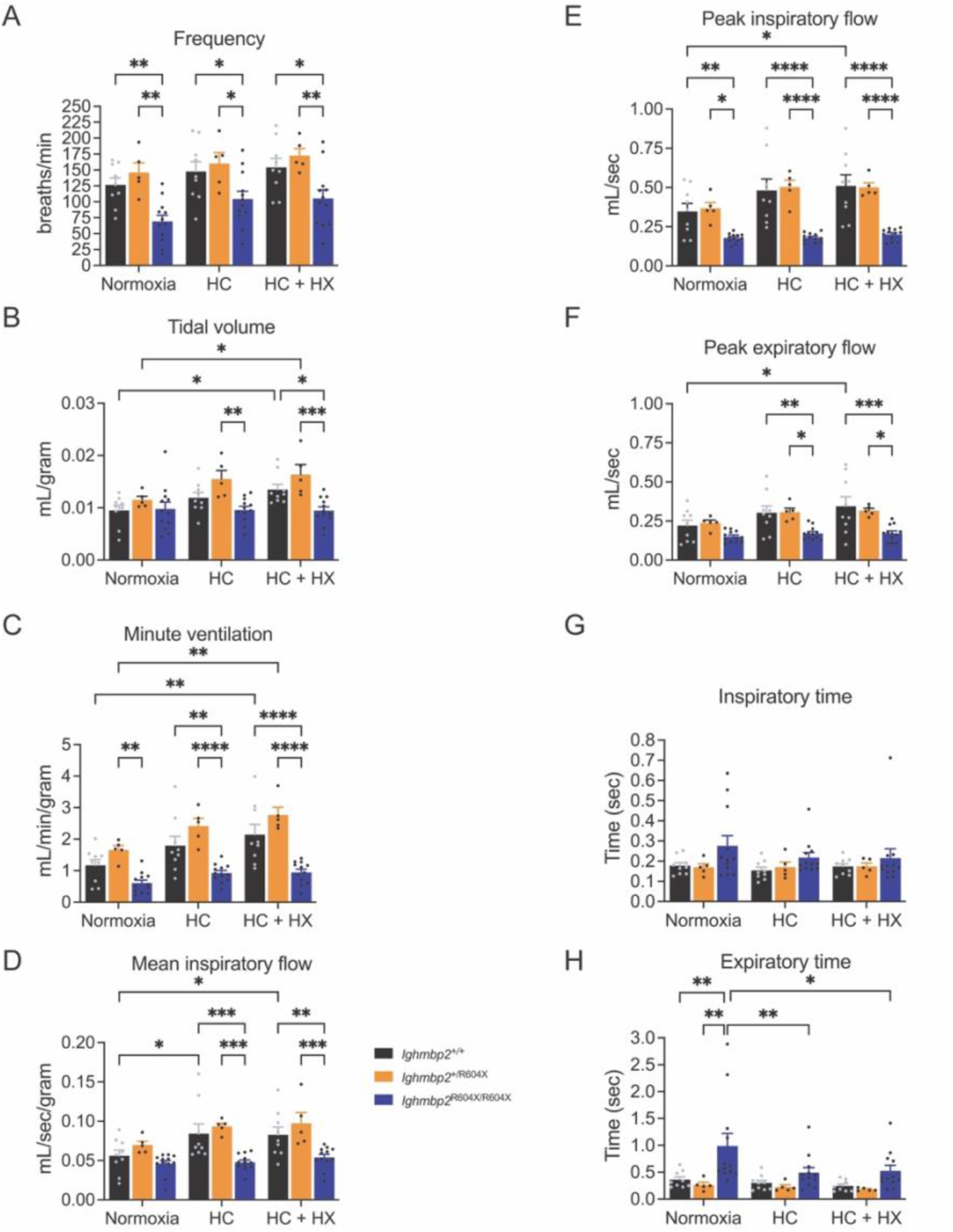
*Ighmbp2*^R604X/R604X^ mice presented with extensive changes in respiration. Wild type (black-9 mice), *Ighmbp2*^+/R604X^ (orange-5 mice), and *Ighmbp2*^R604X/R604X^ (blue-12 mice). Head-out plethysmography was measured at P2 under conditions of normoxia, hypercapnia (HC) and hypercapnia + hypoxia (HC + HX). (**A**) Frequency (breaths/minute) (normoxia *P*=0.0048 +/+ and R604X/R604X, *P*=0.0016 +/R604X and R604X/R604X; HC *P*=0.0428 +/+ and R604X/R604X, *P*=0.0286 +/R604X and R604X/R604X; HC+HX *P*=0.0195 +/+ and R604X/R604X, *P*=0.0064 +/R604X and R604X/R604X). (**B**) Tidal volume (milliliters/gram) (HC *P*=0.0024 +/R604X and R604X/R604X; HC+HX *P*=0.0004 +/R604X and R604X/R604X; *P*=0.0256 +/+ normoxia and HC+HX; *P*=0.0485 +/R604X normoxia and HC+HX). (**C**) Minute ventilation (milliliters/minute/gram) (HC + HX) (normoxia *P*=0.0027 +/R604X and R604X/R604X; HC *P*=0.0015 +/+ and R604X/R604X, *P*<0.0001 +/R604X and R604X/R604X; HC+HX *P*<0.0001 +/+ and R604X/R604X, +/R604X and R604X/R604X). (**D**) Mean inspiratory flow (milliliters/second/gram) (HC *P*=0.0007 +/+ and R604X/R604X, *P*=0.0005 +/R604X and R604X/R604X; HC+HX *P*=0.0088 +/+ and R604X/R604X, *P*=0.0009 +/R604X and R604X/R604X; +/+ *P*=0.0179 normoxia and HC, *P*=0.0267 normoxia and HC+HX). (**E**) Peak inspiratory flow (milliliters/second) (normoxia *P*=0.0067 +/+ and R604X/R604X, *P*=0.0135 +/R604X and R604X/R604X; HC *P*<0.0001 +/+ and R604X/R604X and +/R604X and R604X/R604X; HC+HX *P*<0.0001 +/+ and R604X/R604X and +/R604X and R604X/R604X; +/+ *P*=0.0164 normoxia and HC+HX). (**F**) Peak expiratory flow (milliliters/second) (HC *P*=0.0041 +/+ and R604X/R604X, *P*=0.0171 +/R604X and R604X/R604X; HC+HX *P*=0.0001 +/+ and R604X/R604X, *P*=0.0102 +/R604X and R604X/R604X; +/+ *P*=0.0141 normoxia and HC+HX). (**G**) Inspiratory time (seconds). (**H**) Expiratory time (seconds) (normoxia *P*=0.0014 +/+ and R604X/R604X, *P*=0.0022 +/R604X and R604X/R604X; R604X/R604X *P*=0.0072 normoxia and HC, *P*=0.0129 normoxia and HC+HX). Statistical analyses: two-way ANOVA with Tukey’s multiple comparisons. Values are expressed as mean. HC=hypercapnia, HC+HX=hypercapnia + hypoxia.

Due to the significant respiratory changes, lung pathology was examined (Figure 4). Thickening of the alveolar wall (cell layer thickness), enhanced injury (tissue injury), presence of a hyaline membrane (acellular deposits in alveolar region) and atelectasis (complete or partial collapse of air spaces) was scored. For each of these pathological examinations, significant differences were observed between wild type and *Ighmbp2*^R604X/R604X^ mice (Figure 4A-D). These results are consistent with the changes observed in inspiratory and expiratory flow, especially during respiratory challenge conditions and suggests lung pathology is an important factor contributing to respiratory complications (Figures 3, 4).

**Figure 4.**
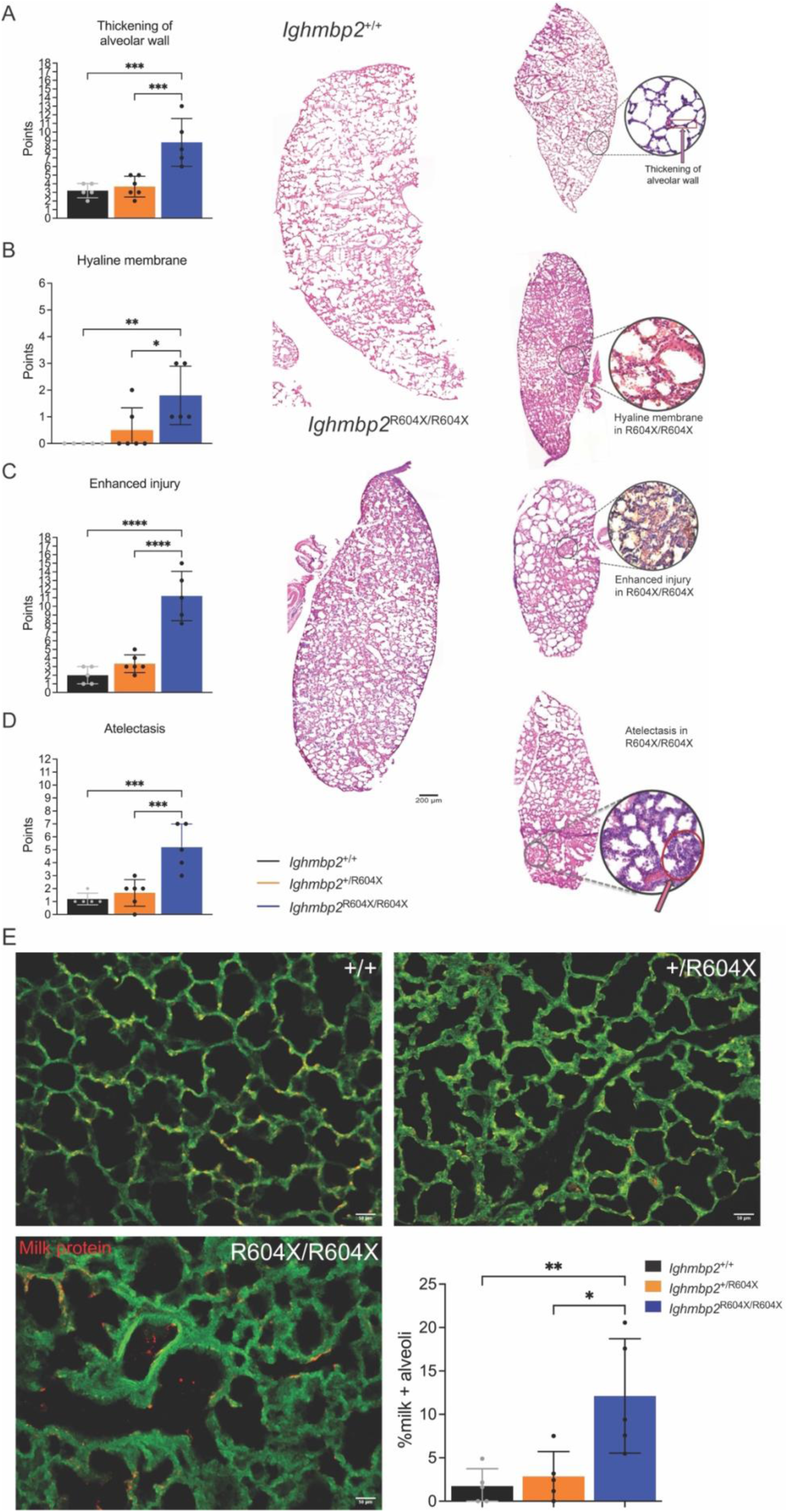
*Ighmbp2*^R604X/R604X^ mice showed changes in lung pathology and evidence of milk aspiration. Wild type (black), *Ighmbp2*^+/R604X^ (orange), and *Ighmbp2*^R604X/R604X^ (blue). Five mice for each genotype and three images per mouse were analyzed. (**A**) Thickening of the alveolar wall was scored 0-18 points. Each image/mouse was scored 0-6 with 0=no thickening of alveolar wall (1 cell layer thick), 1-2=mild thickening, 3-4=moderate thickening of alveolar wall, 5=extensive thickening of alveolar wall, 6=complete thickening. Representative image of thickening of alveolar wall. (**B**) The presence of hyaline membrane (acellular deposits in the alveolar region) was scored 0-6 points. Each image/mouse was scored 0-2 with 0=no acellular debris, 1=partial build up of acellular debris, 2=significant build up of acellular debris. Representative image of hyaline membrane. (**C**) Observed enhanced injury was scored 0-18 points. Each image/mouse was scored 0-6 with 0=no damage, 1=minimum damage, 2=mild damage, 3=moderate damage, 4=pronounced damage, 5=extensive damage, 6=completely damaged. Representative image of enhanced injury in lung. (**D**) Atelectasis (complete or partial collapse of distal air spaces) was scored 0-12. Each image/mouse was scored 0-4 with 0=no collapse, 1=slight collapse, 2=50% collapse, 3=nearly full 75% collapse, 4=full collapse of distal air spaces. Representative image of atelectasis within the lung. (**E**) Representative images of lungs from wild type, *Ighmbp2*^+/R604X^, and *Ighmbp2*^R604X/R604X^ mice lungs immunostained with antibodies against milk proteins (red). The presence of milk within an alveoli was scored based on the presence of anti-milk antibody staining. Five mice were scored per genotype with wild type (708 alveoli scored, 118+ alveoli/mouse), *Ighmbp2*^+/R604X^ (683 alveoli scored, 102+ alveoli/mouse), and *Ighmbp2*^R604X/R604X^ (593 alveoli scored, 91+alveoli/mouse). Wild type (mean=1.73%), *Ighmbp2*^+/R604X^ (mean=2.86%), and *Ighmbp2*^R604X/R604X^ (12.12%) (\**P*=0.0134, \*\**P*=0.0064). Statistical analyses: two-way or one-way ANOVA with Tukey’s multiple comparisons. Values are expressed as mean.

### *Ighmbp2*^R604X/R604X^ mice showed evidence of milk aspiration

*Ighmbp2*^R604X/R604X^ mice showed significant respiratory changes in addition to disease pathology in the lung and reduction in the milk spot; therefore, we examined whether *Ighmbp2*^R604X/R604X^ mice could be aspirating milk into the lungs (Figure 4E). Lungs from wild type, *Ighmbp2*^+/R604X^ and *Ighmbp2*^R604X/R604X^ mice were examined by immunohistochemistry to detect the presence of milk protein. There was significant evidence of milk protein in the lungs of *Ighmbp2*^R604X/R604X^ mice but not in age-matched control mice (mean +/+=1.73%, +/R604X=2.86%, R604X/R604X=12.12%, \**P*=0.0134, \*\**P*=0.0064). These studies provide the first evidence that aspiration could be impacting respiration and survival in *Ighmbp2*^R604X/R604X^ mice.

### Phrenic nerve axons showed reduced size and myelination differences in *Ighmbp2*^R604X/R604X^ mice

The significant respiratory changes in *Ighmbp2*^R604X/R604X^ mice prompted an examination of the phrenic nerve that innervates the diaphragm (Figure 5, Supplemental Figure 1). Phrenic nerve axon area, perimeter, diameter, G-ratio, myelin thickness, number of myelinated fibers/mouse section and distribution were analyzed for wild type, *Ighmbp2*^+/R604X^ and *Ighmbp2*^R604X/R604X^ mice. There was a significant reduction between wild type and *Ighmbp2*^R604X/R604X^ mice in all axon measurements except number of myelinated fibers/mouse section. There were also significant differences between *Ighmbp2*^+/R604X^ and *Ighmbp2*^R604X/R604X^ mice in phrenic nerve axon area, perimeter, diameter and myelin thickness (Figure 5A-C, E, Supplemental Figure 1). When the distribution of phrenic nerve axon area was analyzed, *Ighmbp2*^R604X/R604X^ showed a clear trend towards more, smaller phrenic nerve axons when compared to wild type and *Ighmbp2*^+/R604X^ mice (Figure 5G). Interestingly, while there were no significant respiratory differences between wild type and *Ighmbp2*^+/R604X^ mice observed, phrenic nerve axon perimeter, diameter, G-ratio and myelin thickness were different between these two cohorts (Figure 5B-E). That *Ighmbp2*^R604X/R604X^ mice showed more, smaller phrenic nerve axons with reduced myelin thickness supports the plethysmography findings in these mice.

**Figure 5.**
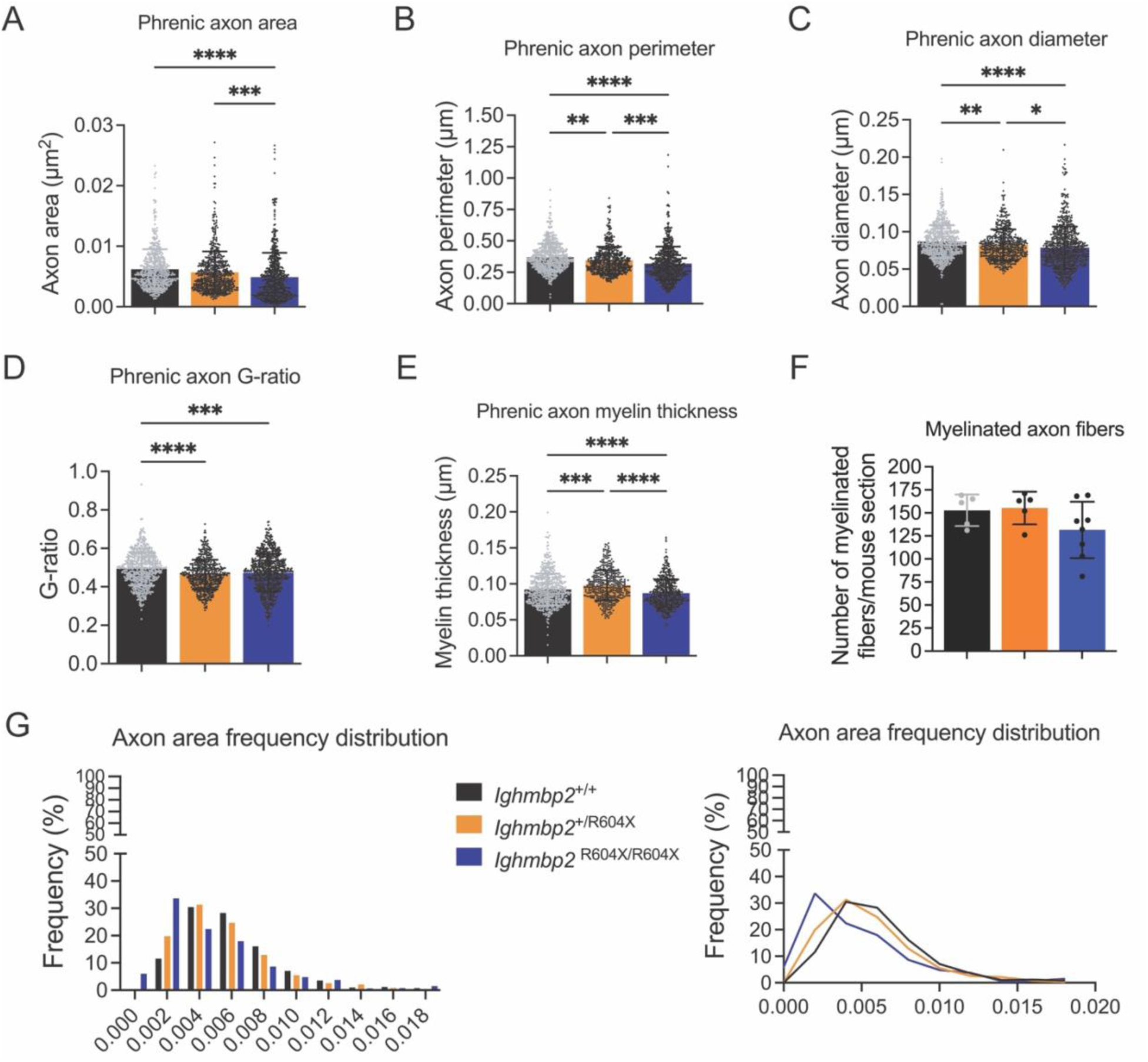
Phrenic nerve axons in *Ighmbp2*^R604X/R604X^ mice were smaller with reduced myelin thickness. Wild type (black), *Ighmbp2*^+/R604X^ (orange), and *Ighmbp2*^R604X/R604X^ (blue). (**A**) Phrenic nerve axon area (+/+=0.0062μm^2^, +/R604X=0.0057μm^2^, R604X/R604X=0.0049μm^2^, *P*<0.0001 +/+ and R604X/R604X, *P*=0.0002 +/R604X and R604X/R604X). (**B**) Phrenic nerve axon perimeter (+/+=0.3693μm, +/R604X=0.3465μm, R604X/R604X=0.3197μm, *P*<0.0001 +/+ and R604X/R604X, *P*=0.0003 +/R604X and R604X/R604X, *P*=0.0057 +/+ and +/R604X). (**C**) Phrenic nerve axon diameter (+/+=0.0868μm, +/R604X=0.0824μm, R604X/R604X=0.0787μm, *P*<0.0001 +/+ and R604X/R604X, *P*=0.0234 +/R604X and R604X/R604X, *P*=0.0090 +/+ and +/R604X). (**D**) Phrenic nerve axon G-ratio (+/+=0.4936, +/R604X=0.4636, R604X/R604X=0.4732, *P*=0.0002 +/+ and R604X/R604X, *P*<0.0001 +/+ and +/R604X). (**E**) Phrenic nerve axon myelin thickness (+/+=0.0923μm, +/R604X=0.0979μm, R604X/R604X=0.0869μm, *P*<0.0001 +/+ and R604X/R604X, *P*<0.0001 +/R604X and R604X/R604X, *P*=0.0001 +/+ and +/R604X). (**F**) Number of myelinated fibers/mouse section (+/+=152.8, +/R604X=155.2, R604X/R604X=131.5). (**G**) Axon area frequency distribution. (**H**) Axon area frequency distribution. Ninety-two to one hundred eleven axons were examined per mouse with five mice per genotype. Statistical analyses: one-way ANOVA with Tukey’s multiple comparisons. Values are expressed as mean.

### Neuromuscular junction innervation of the diaphragm in *Ighmbp2*^R604X/R604X^ mice

The significant respiratory changes and pathology of the phrenic nerve in *Ighmbp2*^R604X/R604X^ mice prompted an examination of the neuromuscular junction (NMJ) innervation status of the diaphragm (Figure 6). Small but significant differences were observed in NMJ innervation of the diaphragm (mean fully innervated endplates +/+=99.5%, +/R604X=98.9%, R604X/R604X=97.4%, *P*=0.0055). *Ighmbp2*^R604X/R604X^ mice did show increased number of NMJs per section suggesting either delayed or absent axon pruning or abnormal sprouting (mean +/+=153.6, +/R604X=144.8, R604X/R604X=198.2, *P*=0.0447) (Figure 6B-E).

**Figure 6.**
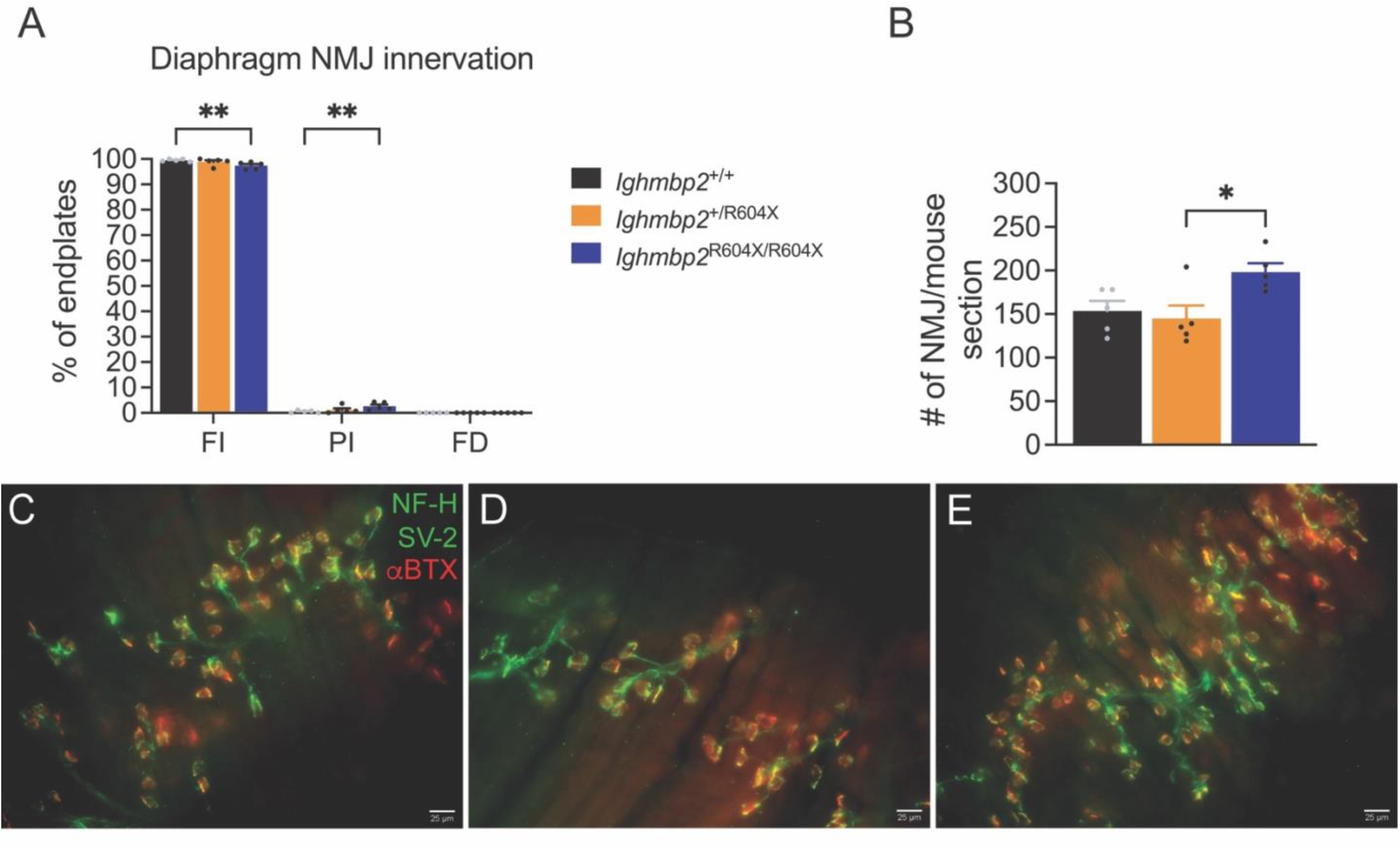
The diaphragm showed slight changes in NMJ innervation in *Ighmbp2*^R604X/R604X^ mice. Wild type (black), *Ighmbp2*^+/R604X^ (orange), and *Ighmbp2*^R604X/R604X^ (blue). (**A**) Diaphragm neuromuscular junction (NMJ) innervation status. The percent of fully innervated endplates (+/+=99.5, +/R604X=98.9, R604X/R604X=97.4, *P*=0.0055). (**B**) Number of diaphragm NMJ/mouse/section (+/+=153.6, +/R604X=144.8, R604X/R604X=198.2, *P*=0.0447). (**C-E**) Representative images of diaphragm NMJ in wild type (**C**), *Ighmbp2*^+/R604X^ (**D**) and *Ighmbp2*^R604X/R604X^ mice (**E**). Green represents neurofilament heavy (NF-H) and synaptic vesicle 2 (SV2) immunostaining while red represents bungarotoxin 594 immunofluorescence (αBTX). Statistical analyses: two-way ANOVA with Tukey’s multiple comparisons. Five mice from each genotype were analyzed with one hundred nineteen to two hundred thirty-three NMJ analyzed/mouse. Values are expressed as mean. FI=fully innervated, PI=partially innervated, FD=fully denervated. NMJ=neuromuscular junction.

### Diaphragm muscle fiber type composition was not altered but muscle fibers were smaller in Ighmbp2^R604X/R604X^ mice

Changes in muscle fiber type composition and the size of muscle fibers could alter the function or the efficiency of the diaphragm. To determine whether any changes in the diaphragm muscle were present, muscle fiber area and muscle fiber type composition were examined in wild type, *Ighmbp2*^+/R604X^ and *Ighmbp2*^R604X/R604X^ mice (Figure 7, Supplemental Figure 2). Skeletal muscle is comprised of distinct myofibers with classifications based on expression of different myosin heavy chains (MyHC). MyHC-emb encoded by *Myh3* is expressed embryonic day 9.5 to 13.5 in mice and is not normally expressed in adult muscles except transiently during skeletal muscle regeneration. Fibers expressing MyHC type 1 are classified as slow twitch fibers, present an oxidative metabolic type and express *Myh7*. MyHC type 2 fibers are classified as fast twitch and are comprised of MyHC 2A, MyHC 2X, and MyHC 2B myofibers that express *Myh2*, *Myh1*, and *Myh4*, respectively. MyHC 2A is oxidative in metabolism while 2X and 2B are glycolytic (29–31). When diaphragm muscle fiber area was quantified, *Ighmbp2*^R604X/R604X^ mice muscle fibers were significantly smaller and the distribution demonstrated that *Ighmbp2*^R604X/R604X^ mice had more, small diaphragm muscle fibers than either wild type or *Ighmbp2*^+/R604X^ mice (Figure 7A, J). Diaphragm embryonic muscle fibers had smaller area in *Ighmbp2*^+/R604X^ and *Ighmbp2*^R604X/R604X^ mice when compared to wild type mice and while there were more embryonic muscle fibers in *Ighmbp2*^R604X/R604X^ mice it was not significant (Figure 7B, K). Regardless of slow or fast twitch fibers or metabolism, *Ighmbp2*^R604X/R604X^ mice had significantly reduced diaphragm muscle fiber area; however, there were no large differences between wild type and *Ighmbp2*^R604X/R604X^ mice in the composition of the diaphragm muscle (Figure 7B-I, K). In *Ighmbp2*^R604X/R604X^ mice, the largest percentage of diaphragm muscle fiber type was MyHC-emb muscle fibers (emb=65.5%, Type 1=6.3%, Type 2A=15.3%, Type 2B=11.6%, unlabelled=31.4%) and MyHC-emb fibers also saw the most dramatic reduction in size (49%) (Figure 7B, K). These results show that *Ighmbp2*^R604X/R604X^ mice have smaller diaphragm muscle fibers of all MyHC types and suggest diaphragm function could be impacted by these changes, consistent with the plethysmography data. As well, a large portion of *Ighmbp2*^R604X/R604X^ diaphragm muscle fibers were MyHC-emb suggesting that there was substantial regeneration of muscle fibers occurring in *Ighmbp2*^R604X/R604X^ mice.

**Figure 7.**
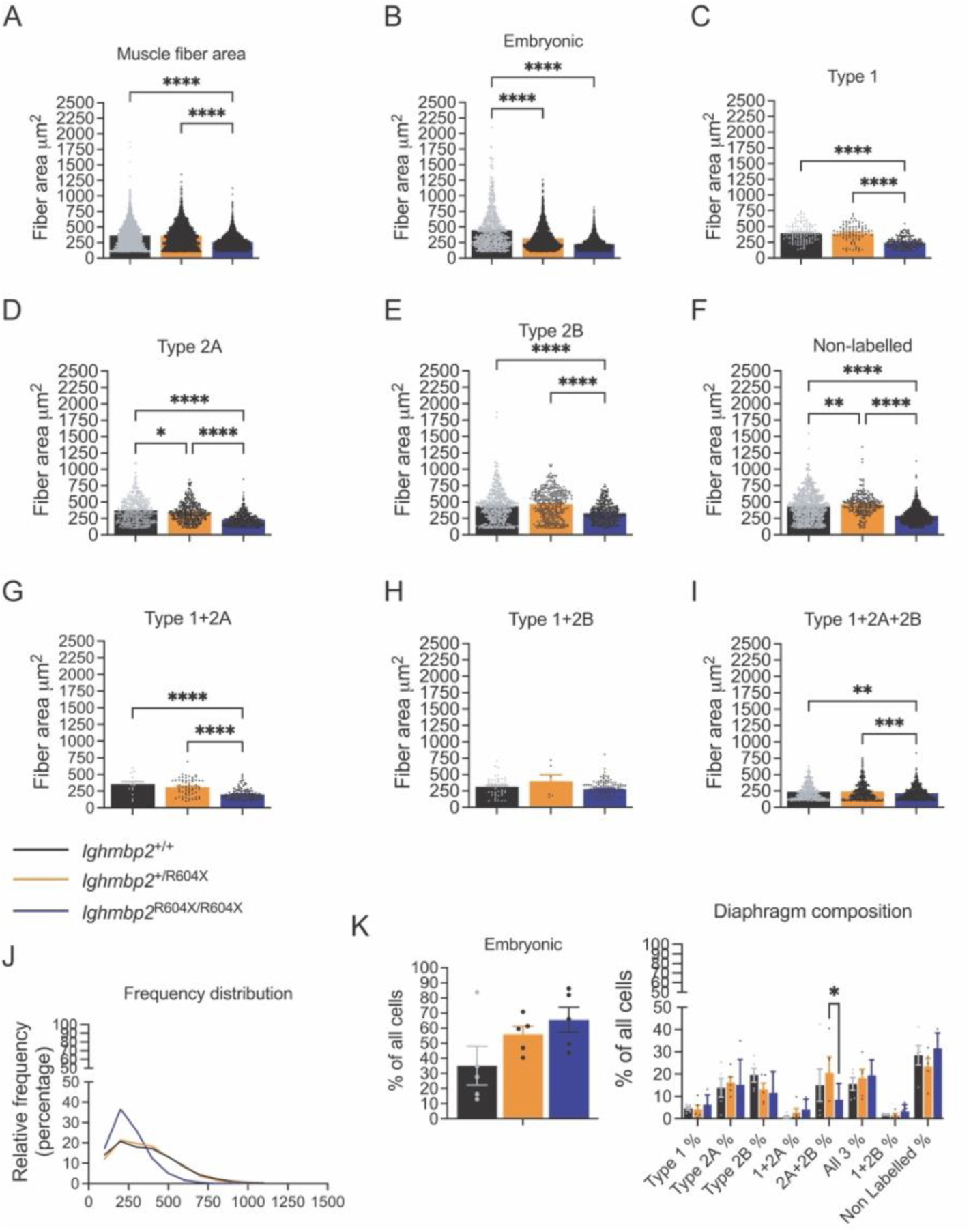
Diaphragm muscle fiber type composition was not altered but muscle fibers were smaller in *Ighmbp2*^R604X/R604X^ mice. Wild type (black), *Ighmbp2*^+/R604X^ (orange), and *Ighmbp2*^R604X/R604X^ (blue). (**A**) Diaphragm muscle fiber area (+/+=364.7μm^2^, +/R604X=358.3μm^2^, R604X/R604X=261.8μm^2^, *P*<0.0001 +/+ and R604X/R604X and +/R604X and R604X/R604X). (**B**) Diaphragm embryonic muscle fiber area (+/+=449.3μm^2^, +/R604X=317.7μm^2^, R604X/R604X=227.7μm^2^, *P*<0.0001 +/+ and R604X/R604X and +/+ and +/R604X). (**C**) Diaphragm Type 1 muscle fiber area (+/+=396μm^2^, +/R604X=390.7μm^2^, R604X/R604X=242.4μm^2^, *P*<0.0001 +/+ and R604X/R604X and +/R604X and R604X/R604X). (**D**) Diaphragm Type 2A muscle fiber area (+/+=373.5μm^2^, +/R604X=347.3μm^2^, R604X/R604X=231.2μm^2^, *P*<0.0001 +/+ and R604X/R604X and +/R604X and R604X/R604X, *P*=0.0275 +/+ and +/R604X). (**E**) Diaphragm Type 2B muscle fiber area (+/+=437.3μm^2^, +/R604X=464.7μm^2^, R604X/R604X=329.4μm^2^, *P*<0.0001 +/+ and R604X/R604X and +/R604X and R604X/R604X). (**F**) Diaphragm non-labelled (representing embryonic and Type X) muscle fiber area (+/+=430.7μm^2^, +/R604X=467.9μm^2^, R604X/R604X=290.3μm^2^, *P*<0.0001 +/+ and R604X/R604X and +/R604X and R604X/R604X, *P*=0.0062 +/+ and +/R604X). (**G**) Diaphragm Type 1+2A hybrid muscle fiber area (+/+=352.4μm^2^, +/R604X=308.7μm^2^, R604X/R604X=201.4μm^2^, *P*<0.0001 +/+ and R604X/R604X and +/R604X and R604X/R604X). (**H**) Diaphragm Type 1+2B hybrid muscle fiber area (+/+=316.9μm^2^, +/R604X=394.7μm^2^, R604X/R604X=280.2μm^2^). (**I**) Diaphragm Type 1+2A+2B muscle fiber area (+/+=237.9μm^2^, +/R604X=242.4μm^2^, R604X/R604X=215.1μm^2^, *P*=0.0060 +/+ and R604X/R604X, *P*=0.0003 +/R604X and R604X/R604X). (**J**) Frequency distribution of diaphragm muscle fiber area in wild type, *Ighmbp2*^+/R604X^ and *Ighmbp2*^R604X/R604X^ mice. (**K**) Percentage of all cells expressing a given muscle fiber type in wild type, *Ighmbp2*^+/R604X^ and *Ighmbp2*^R604X/R604X^ mice, *P*=0.0491. Five mice from each genotype were analyzed with 2434-3177 total muscle fibers analyzed per genotype. Statistical analyses: one-way ANOVA with Dunnett’s multiple comparisons. Values are expressed as mean.

### The sciatic nerve in *Ighmbp2*^R604X/R604X^ mice showed deficits in size and myelination

Since the phrenic nerve showed significant changes in axon size and myelination we also examined the same measurements in the sciatic nerve that innervates hindlimb muscles to determine if there was similar pathology (Figure 8, Supplemental Figure 1). Consistent with the phrenic nerve, there was significant reduction in sciatic nerve axon area, perimeter, diameter, G-ratio and myelin thickness between *Ighmbp2*^R604X/R604X^ and wild type and *Ighmbp2*^+/R604X^ mice (Figure 8A-H). *Ighmbp2*^+/R604X^ mice demonstrated differences in sciatic nerve axon perimeter, diameter and G-ratio when compared to wild type mice, consistent with the data observed in the phrenic axons (Figure 8). There was also a reduction in the number of myelinated axons scored per mouse section in *Ighmbp2*^R604X/R604X^ mice (Figure 8F). When the distribution of sciatic nerve axon area was examined *Ighmbp2*^R604X/R604X^ mice had more, small axon fibers when compared to *Ighmbp2*^+/R604X^ or wild type mice (0.004 or less, +/+=42%, +/R604X=50%, R604X/R604X=70%) (Figure 8G, H).

**Figure 8.**
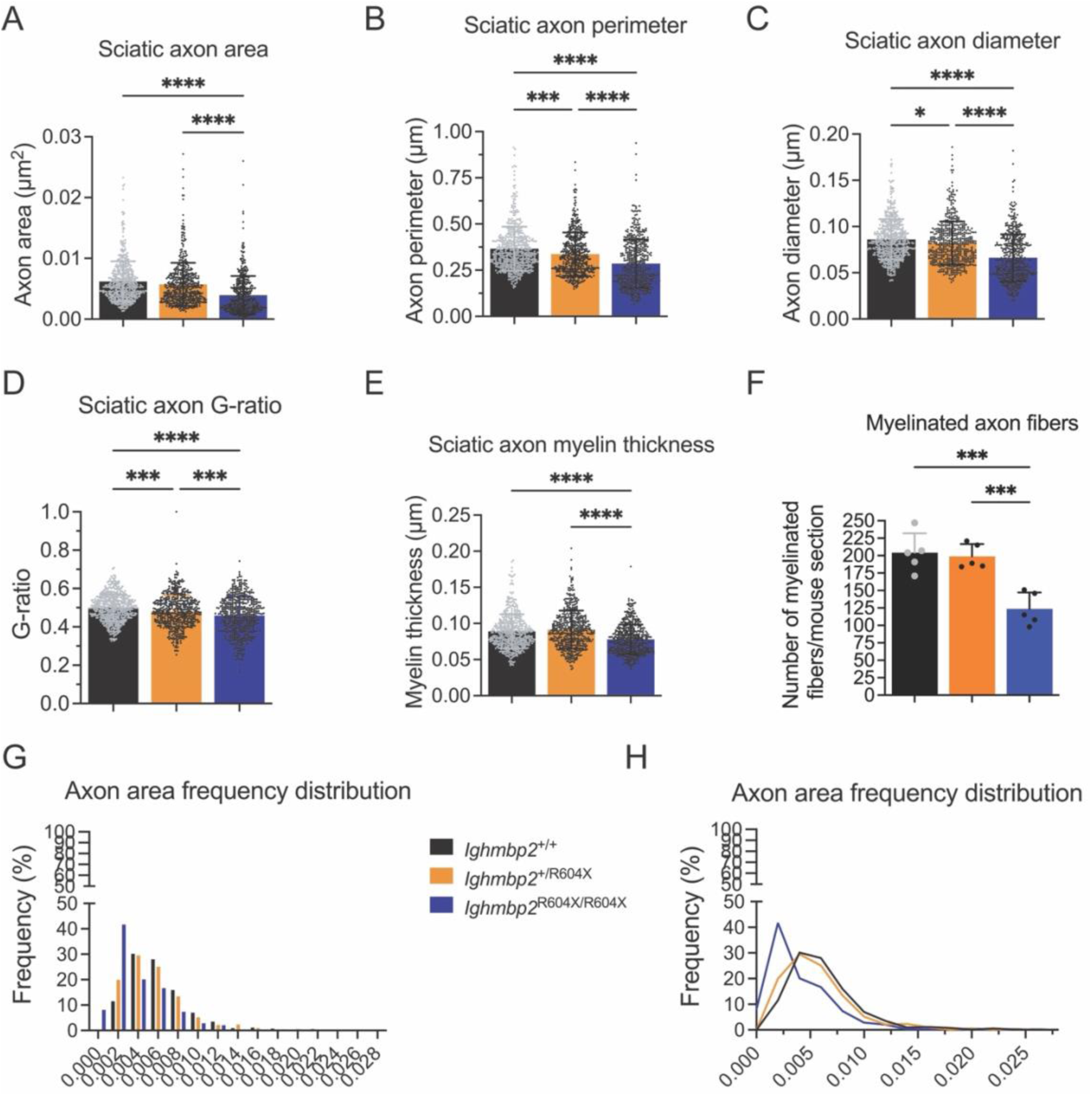
The sciatic nerve in *Ighmbp2*^R604X/R604X^ mice showed changes in size and myelination. Wild type (black), *Ighmbp2*^+/R604X^ (orange), and *Ighmbp2*^R604X/R604X^ (blue). (**A**) Sciatic nerve axon area (+/+=0.006μm^2^, +/R604X=0.006μm^2^, R604X/R604X=0.004μm^2^, *P*<0.0001 +/+ and R604X/R604X and +/R604X and R604X/R604X). (**B**) Sciatic nerve axon perimeter (+/+=0.366μm, +/R604X=0.337μm, R604X/R604X=0.285μm, *P*<0.0001 +/+ and R604X/R604X and +/R604X and R604X/R604X, *P*=0.0004 +/+ and +/R604X). (**C**) Sciatic nerve axon diameter (+/+=0.086μm, +/R604X=0.082μm, R604X/R604X=0.066μm, *P*<0.0001 +/+ and R604X/R604X and +/R604X and R604X/R604X, *P*=0.0173 +/+ and +/R604X). (**D**) Sciatic nerve axon G-ratio (+/+=0.500, +/R604X=0.480, R604X/R604X=0.456, *P*<0.0001 +/+ and R604X/R604X, *P*=0.0002 +/R604X and R604X/R604X, P=0.0008 +/+ and +/R604X). (**E**) Sciatic nerve axon myelin thickness (+/+=0.089μm, +/R604X=0.092μm, R604X/R604X=0.078μm, *P*<0.0001 +/+ and R604X/R604X and +/R604X and R604X/R604X). (**F**) Number of myelinated fibers/mouse section (+/+=204, +/R604X=199, R604X/R604X=124, *P*=0.0004 +/+ and R604X/R604X, *P*=0.0007 +/R604X and R604X/R604X). (**G**) Axon area frequency distribution, percentage of area 0-0.004μm^2^ +/+=42%, +/R604X=50% and R604X/R604X=70%. (**H**) Axon area frequency distribution. Five mice from each genotype were analyzed with ninety-three to one hundred six axons analyzed per mouse. Statistical analyses: one-way ANOVA with Tukey’s multiple comparisons. Values are expressed as mean.

### Neuromuscular junction innervation of the forelimb and hindlimb muscles was altered in Ighmbp^2R604X/R604X^ mice

The average lifespan of *Ighmbp2*^R604X/R604X^ mice was six days. To determine whether there was significant forelimb and hindlimb pathology by P6, we examined NMJ innervation and muscle composition in forelimb (tricep) or hindlimb (gastrocnemius (gastroc), tibialis anterior (TA)) muscles (Figure 9, Supplemental Figure 3). Significant differences were observed in NMJ innervation of the TA muscle of *Ighmbp2*^R604X/R604X^ mice (mean fully innervated endplates +/+=99.4%, +/R604X=98.3%, R604X/R604X=40.8%, *P*<0.0001) (mean fully denervated endplates +/+=0.3%, +/R604X=0.4%, R604X/R604X=36.0%, *P*<0.0001) (Figure 9A, D). Significant NMJ denervation of the hindlimb gastrocnemius muscle was also observed (mean fully innervated endplates +/+=98.0%, +/R604X=98.6%, R604X/R604X=49.7%, *P*<0.0001) (mean fully denervated endplates +/+=1.2%, +/R604X=0.5%, R604X/R604X=31.2%, *P*<0.0001) (Figure 9B, D). Interestingly, the forelimb tricep muscle showed less NMJ denervation (mean fully innervated endplates +/+=90.5%, +/R604X=96.3%, R604X/R604X=69.2%, \*\**P*=0.0019, \*\*\**P*=0.0001) (Figure 9C, Supplmental Figure 3). These results suggest that forelimb and hindlimb motor function would be impacted by the significant NMJ denervation and is consistent with the sciatic nerve pathology.

**Figure 9.**
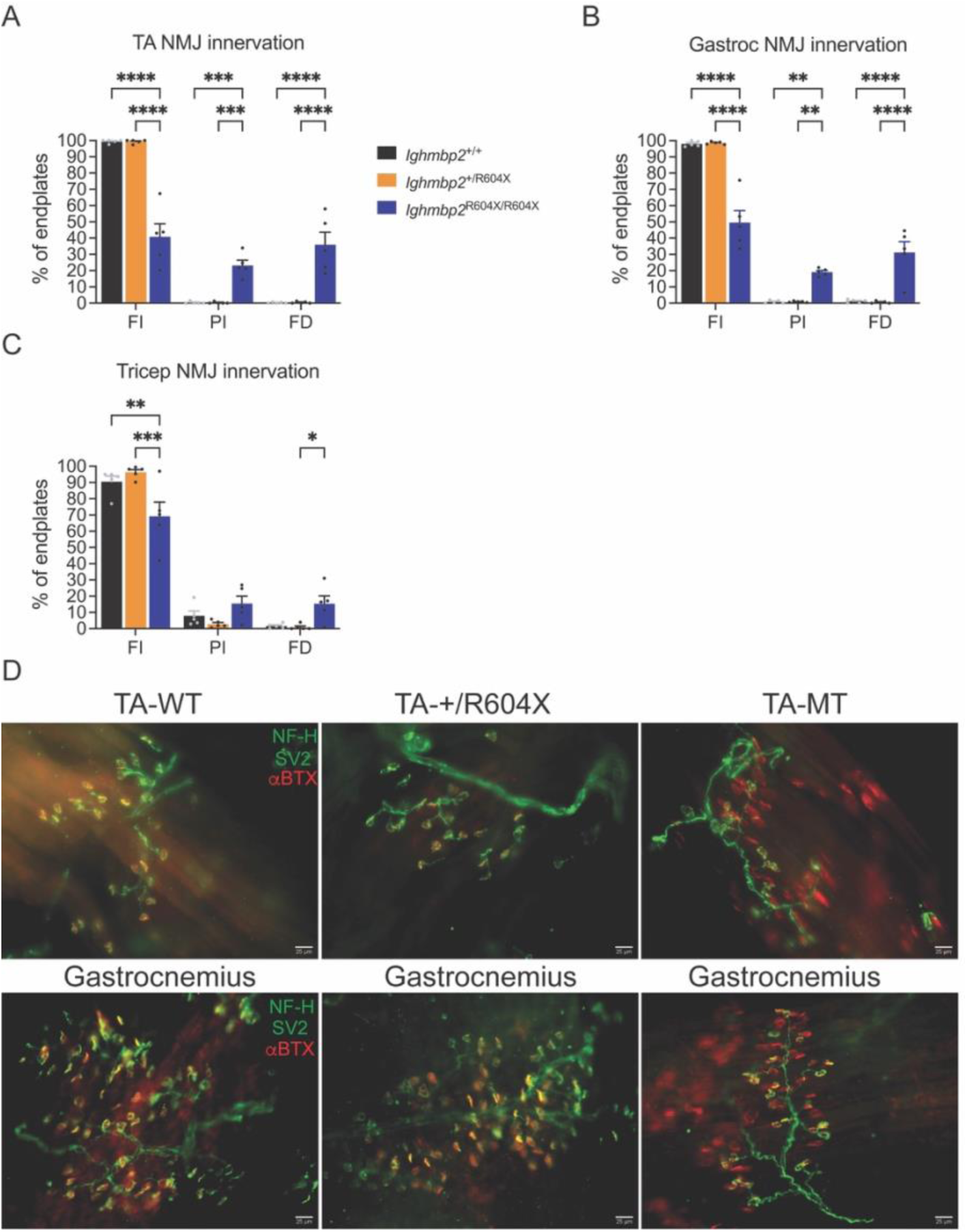
Hindlimb muscles TA and gastrocnemius and to an extent the forelimb tricep showed NMJ denervation in *Ighmbp2*^R604X/R604X^ mice. Wild type (black), *Ighmbp2*^+/R604X^ (orange), and *Ighmbp2*^R604X/R604X^ (blue). (**A**) NMJ innervation status of hindlimb muscle tibialis anterior (TA). The percent of fully innervated endplates (+/+=99.4, +/R604X=99.3, R604X/R604X=40.8, *P*<0.0001). The percent of partially innervated endplates (+/+=0.3, +/R604X=0.3, R604X/R604X=23.2, *P*=0.0004-0.0005). The percent of fully denervated endplates (+/+=0.3, +/R604X=0.3, R604X/R604X=36.0, *P*<0.0001). (**B**) NMJ innervation status of hindlimb muscle gastrocnemius. The percent of fully innervated endplates (+/+=98.0, +/R604X=99.6, R604X/R604X=49.7, *P*<0.0001). The percent of partially innervated endplates (+/+=0.8, +/R604X=0.9, R604X/R604X=19.1, *P*=0.0012-0.0013). The percent of fully denervated endplates (+/+=1.2, +/R604X=0.5, R604X/R604X=31.2, *P*<0.0001). (**C**) NMJ innervation status of forelimb muscle tricep. The percent of fully innervated endplates (+/+=90.5, +/R604X=96.3, R604X/R604X=69.2, *P*=0.0010 and *P*=0.0001, respectively). The percent of partially innervated endplates (+/+=7.9, +/R604X=2.8, R604X/R604X=15.5). The percent of fully denervated endplates (+/+=1.6, +/R604X=0.9, R604X/R604X=15.3, *P*=0.0425). (**D**) Representative images of neuromuscular junctions (NMJ) of the TA and gastrocnemius. Green represents neurofilament heavy (NF-H) and synaptic vesicle 2 (SV2) immunostaining while red represents bungarotoxin 594 immunofluorescence (αBTX). Statistical analyses: two-way ANOVA with Tukey’s multiple comparisons. Five mice from each genotype were analyzed with ninety-three to three hundred sixty-nine NMJ counted per mouse. Values are expressed as mean. FI=fully innervated, PI=partially innervated, FD=fully denervated, NMJ=neuromuscular junction, WT= wild type, MT=R604X/R604X, TA=tibialis anterior.

### *Ighmbp2*^R604X/R604X^ forelimb and hindlimb muscles showed significant atrophy with changes in muscle fiber composition

To determine the extent that forelimb and hindlimb muscles showed disease pathology in *Ighmbp2*^R604X/R604X^ mice, the tricep, tibialis anterior (TA) and gastrocnemius muscles were examined (Figures 10-12, Supplemental Figures 2, 4). Significant NMJ denervation was observed in *Ighmbp2*^R604X/R604X^ mice with the hindlimb muscles changed more than tricep (Figure 9). In *Ighmbp2*^R604X/R604X^ mice, there was significant reduction in TA muscle fiber area when compared to wild type and *Ighmbp2*^+/R604X^ mice (Figure 10A). When TA muscle fiber types were examined in *Ighmbp2*^R604X/R604X^ mice, there were significant reductions in area of all muscle fiber types except embryonic muscle fibers that were larger (Figure 10B-F). Interestingly, in *Ighmbp2*^+/R604X^ mice TA embryonic, Type IIB and non-labelled muscle fibers were also statistically different in size from wild type mice (Figure 10B, E, F). When the overall TA muscle fiber type composition was analyzed, *Ighmbp2*^R604X/R604X^ mice demonstrated increased percentage of TA embryonic muscle fibers when compared to wild type and *Ighmbp2*^+/R604X^ mice (mean +/+=32.2%, +/R604X=34.7%, R604X/R604X=65.6%, *P*=0.0002 and *P*=0.0005) suggesting the TA was undergoing muscle regeneration. Additionally, there were significant changes in TA muscle Type 2B composition in *Ighmbp2*^R604X/R604X^ mice (mean +/+=45.4%, +/R604X=27.2%, R604X/R604X=7.8%, *P*<0.0001) showing a reduction of the faster and larger muscle fiber types (Figure 10G). These shifts suggest a delay in maturation to large, fast twitch glycolytic muscle fibers. The increased non-labelled composition in *Ighmbp2*^R604X/R604X^ mice (representing embryonic and Type 2X fibers) likely reflects increased embryonic muscle fibers (mean +/+=35.8%, +/R604X=54.7%, R604X/R604X=69.3%, *P*<0.0001).

**Figure 10.**
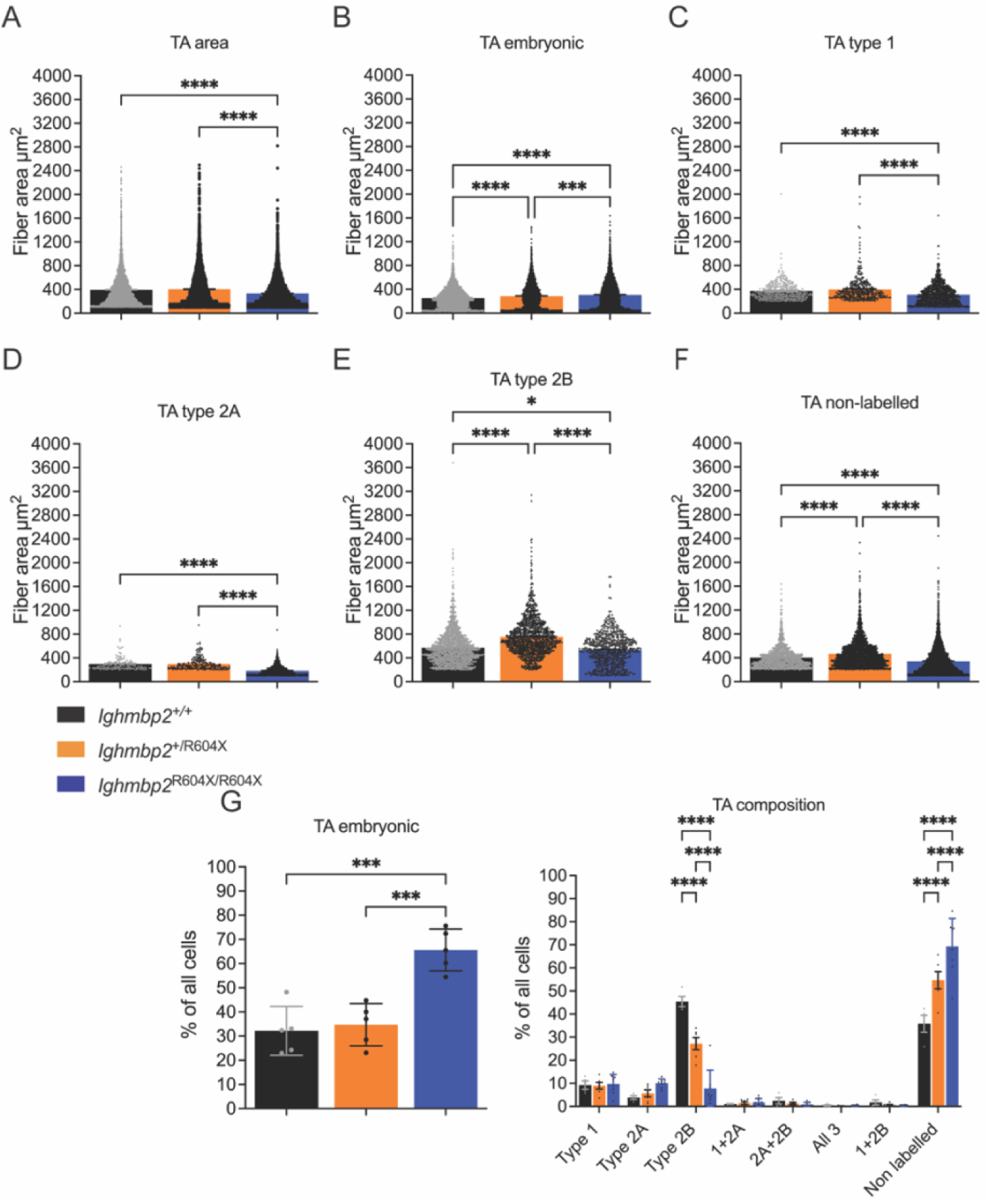
TA muscle fiber muscle fiber size and types were altered in *Ighmbp2*^R604X/R604X^ mice. Wild type (black), *Ighmbp2*^+/R604X^ (orange), and *Ighmbp2*^R604X/R604X^ (blue). (**A**) TA total muscle fiber area (+/+=393.3μm^2^, +/R604X=403.8μm^2^, R604X/R604X=331.5μm^2^, *P*<0.0001). (**B**) TA embryonic muscle fiber area (+/+=252.5μm^2^, +/R604X=285.7μm^2^, R604X/R604X=305.9μm^2^, *P*<0.0001 +/+ and R604X/R604X, *P*<0.0001 +/+ and +/R604X, *P*=0.0001 +/R604X and R604X/R604X). (**C**) TA Type 1 muscle fiber area (+/+=372.5μm^2^, +/R604X=399.6μm^2^, R604X/R604X=310.8μm^2^, *P*<0.0001). (**D**) TA Type 2A muscle fiber area (+/+=296.3μm^2^, +/R604X=297.8μm^2^, R604X/R604X=183.3μm^2^, *P*<0.0001). (**E**) TA Type 2B muscle fiber area (+/+=570.3μm^2^, +/R604X=753.4μm^2^, R604X/R604X=534.0μm^2^, *P*<0.0001 +/+ and +/R604X and +/R604X and R604X/R604X, *P*=0.0252 +/+ and R604X/R604X). (**F**) TA non-labelled (representing embryonic and Type X) muscle fiber area (+/+=402.0μm^2^, +/R604X=468.5μm^2^, R604X/R604X=340.5μm^2^, *P*<0.0001). (**G**) Percentage of all cells expressing a given muscle fiber type in wild type, *Ighmbp2*^+/R604X^ and *Ighmbp2*^R604X/R604X^ mice (embryonic +/+=32.2, +/R604X=34.7, R604X/R604X=65.6, *P*=0.0002 +/+ and R604X/R604X, *P*=0.0005 +/R604X and R604X/R604X) (Type 2B +/+=45.4, +/R604X=27.2, R604X/R604X=7.8, *P*<0.0001) (non-labelled +/+=35.8, +/R604X=54.7, R604X/R604X=69.3, *P*<0.0001). Five mice from each genotype were analyzed. Statistical analyses: one-way ANOVA with Dunnett’s multiple comparisons. Values are expressed as mean.

When the gastrocnemius muscle was examined in wild type, *Ighmbp2*^+/R604X^ and *Ighmbp2*^R604X/R604X^ mice, there were significant reductions in muscle fiber area across all fiber types except Type 1 where there were no differences between the genotypes (Figure 11, Supplemental Figure 4). Similar to the TA, there were differences between wild type and *Ighmbp2*^+/R604X^ mice in muscle fiber type area except in Type 1 fibers. Interestingly, *Ighmbp2*^+/R604X^ and *Ighmbp2*^R604X/R604X^ mice showed similar area in embryonic, Type 1 and Type 2A muscle fibers (Figure 11A-D). When the overall gastrocnemius muscle fiber type composition was analyzed, unlike the TA, *Ighmbp2*^R604X/R604X^ mice showed no differences from wild type mice in embryonic muscle fiber type (mean +/+=62.5%, +/R604X=47.1%, R604X/R604X=58.9%) (Figure 11G). There were however, more non-labelled fibers in *Ighmbp2*^R604X/R604X^ mice (mean +/+=23.7%, +/R604X=29.6%, R604X/R604X=49.4%).

**Figure 11.**
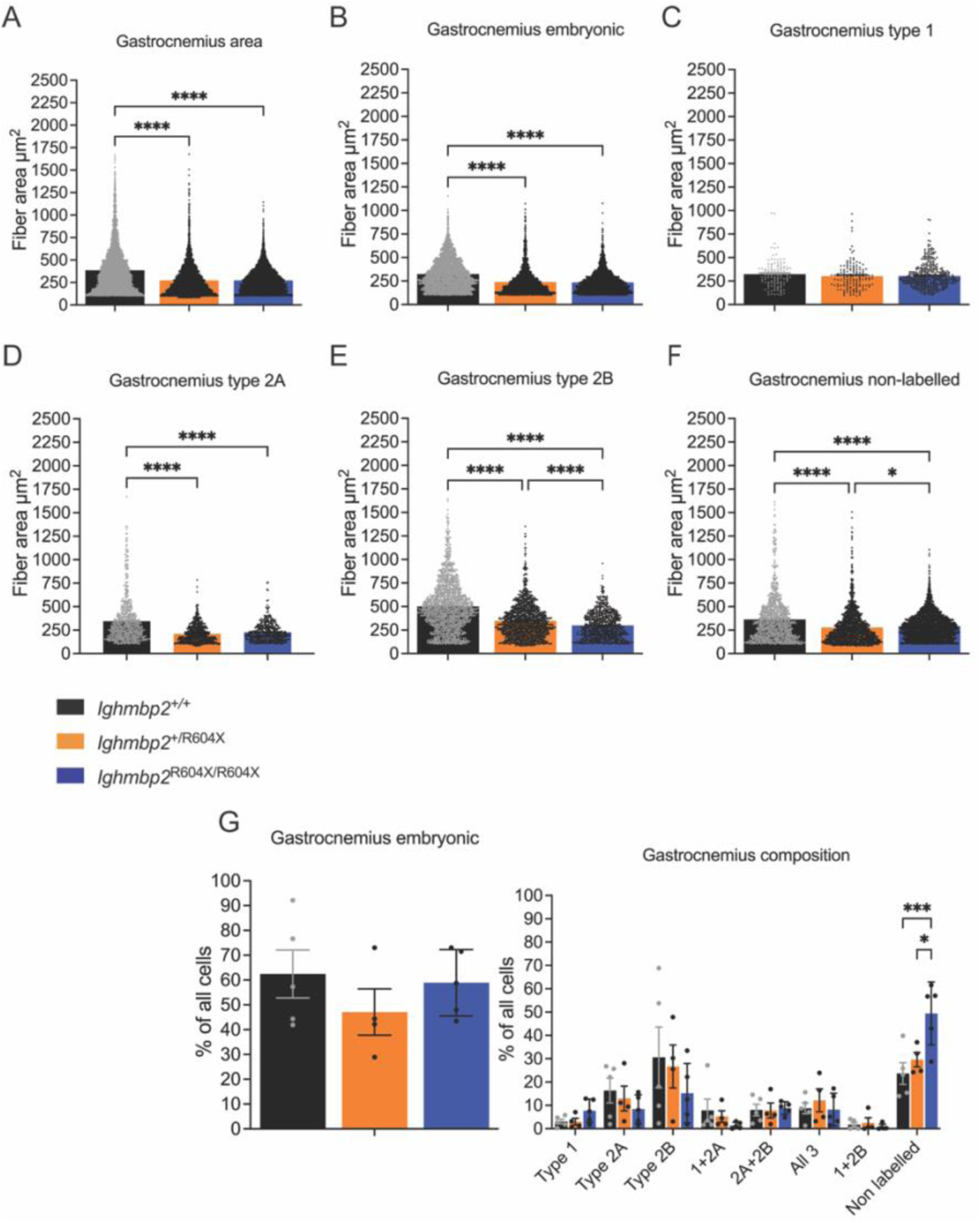
*Ighmbp2*^R604X/R604X^ mice showed reduced gastrocnemius muscle fiber size in all fibers but type I. Wild type (black), *Ighmbp2*^+/R604X^ (orange), and *Ighmbp2*^R604X/R604X^ (blue). (**A**) Gastrocnemius muscle fiber area (+/+=387.5m^2^, +/R604X=273.3μm^2^, R604X/R604X=273.3μm^2^, *P*<0.0001). (**B**) Gastrocnemius embryonic muscle fiber area (+/+=323.8μm^2^, +/R604X=240.8μm^2^, R604X/R604X=235.6μm^2^, *P*<0.0001). (**C**) Gastrocnemius Type 1 muscle fiber area (+/+=322.7μm^2^, +/R604X=301.8μm^2^, R604X/R604X=305.2μm^2^). (**D**) Gastrocnemius Type 2A muscle fiber area (+/+=342.1μm^2^, +/R604X=211.0μm^2^, R604X/R604X=227.1μm^2^, *P*<0.0001). (**E**) Gastrocnemius Type 2B muscle fiber area (+/+=500.8μm^2^, +/R604X=347.7μm^2^, R604X/R604X=298.8μm^2^, *P*<0.0001). (**F**) Gastrocnemius non-labelled (representing embryonic and Type X) muscle fiber area (+/+=363.4μm^2^, +/R604X=276.9μm^2^, R604X/R604X=292.4μm^2^, *P*<0.0001, *P*=0.0155 +/R604X and R604X/R604X). (**G**) Percentage of all cells expressing a given muscle fiber type in wild type, *Ighmbp2*^+/R604X^ and *Ighmbp2*^R604X/R604X^ mice (embryonic +/+=62.5, +/R604X=47.1, R604X/R604X=58.9) (non-labelled +/+=23.7, +/R604X=29.6, R604X/R604X=49.4, *P*=0.0004 +/+ and R604X/R604X, *P*=0.0133 +/R604X and R604X/R604X). Five mice from each genotype were analyzed. Statistical analyses: one-way ANOVA with Dunnett’s multiple comparisons. Values are expressed as mean.

When tricep muscle fiber types were examined, there was a significant reduction in *Ighmbp2*^R604X/R604X^ muscle fiber area found in all muscle fiber types except Type 2A when compared to wild type mice (Figure 12, Supplemental Figure 4). As well, there were clear differences in muscle fiber type area between wild type and *Ighmbp2*^+/R604X^ mice (Figure 12B-F). Interestingly, unlike the diaphragm, there were differences in muscle fiber type composition between wild type and *Ighmbp2*^R604X/R604X^ mice in embryonic, Type 2B and non-labelled fibers with a significant reduction in Type 2B fibers and a significant increase in embryonic and non-labelled fibers (R604X/R604X emb=34.6%, Type 1=6.7%, Type 2A=5.0%, Type 2B=1.6%, non-labelled=85.6%) (Figure 12G). This is similar to changes observed in TA muscle fiber composition. The increased embryonic fibers suggest regeneration is occurring while the significant shift from Type 2B suggests a delay to large, fast twitch glycolytic muscle fibers.

**Figure 12.**
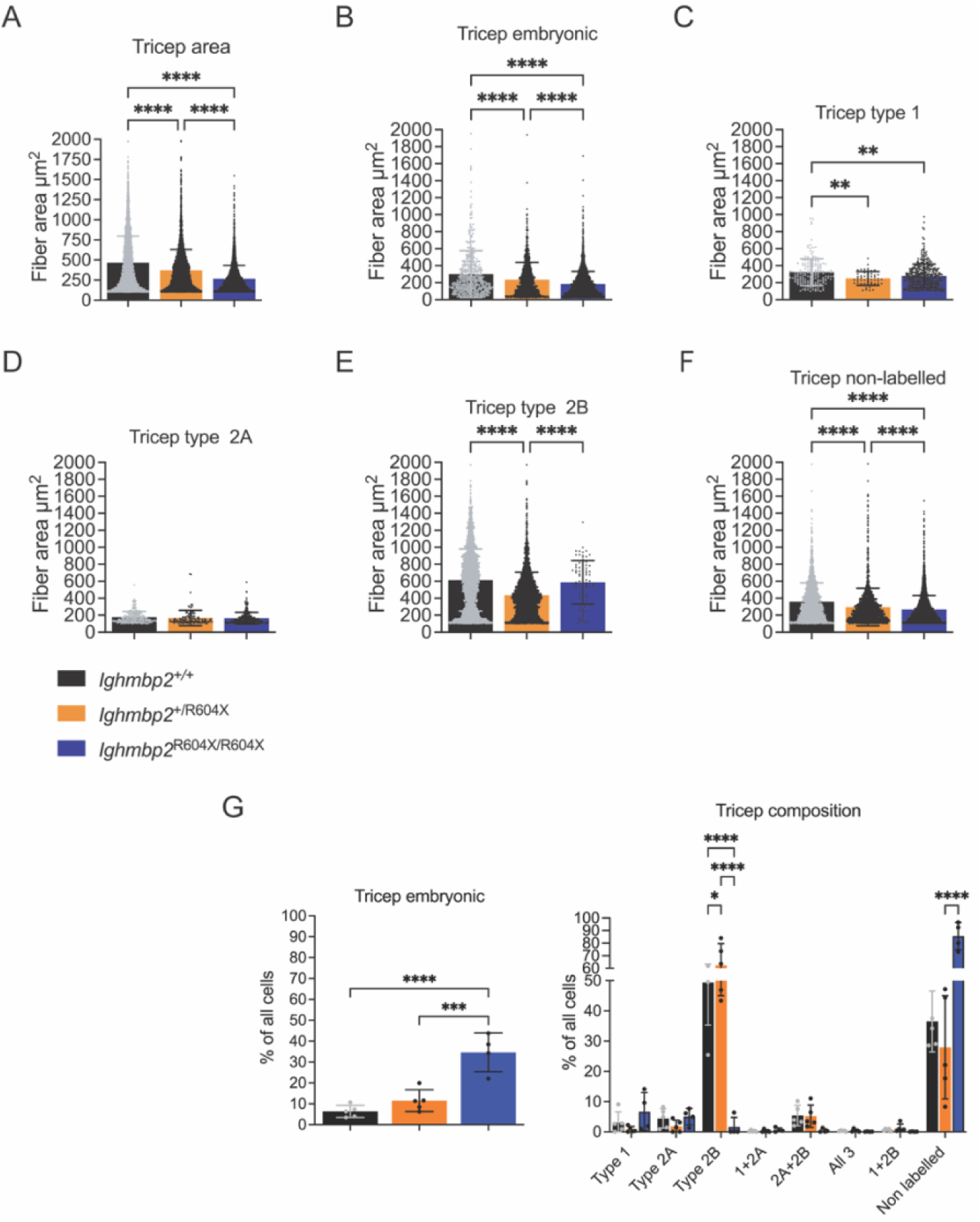
Tricep muscle fiber size and composition were altered in *Ighmbp2*^R604X/R604X^ mice. Wild type (black), *Ighmbp2*^+/R604X^ (orange), and *Ighmbp2*^R604X/R604X^ (blue). (**A**) Tricep muscle fiber area (+/+=465.2μm^2^, +/R604X=372.5μm^2^, R604X/R604X=267.2μm^2^, *P*<0.0001). (**B**) Tricep embryonic muscle fiber area (+/+=301.4μm^2^, +/R604X=234.9μm^2^, R604X/R604X=184.1μm^2^, *P*<0.0001). (**C**) Tricep Type 1 muscle fiber area (+/+=323.5μm^2^, +/R604X=250.1μm^2^, R604X/R604X=279.7μm^2^, *P*=0.0018 +/+ and R604X/R604X and P=0.0023 +/+ and +/R604X). (**D**) Tricep Type 2A muscle fiber area (+/+=177.4μm^2^, +/R604X=167.6μm^2^, R604X/R604X=164.0μm^2^). (**E**) Tricep Type 2B muscle fiber area (+/+=614.3μm^2^, +/R604X=435.7μm^2^, R604X/R604X=588.0μm^2^, *P*<0.0001 +/+ and +/R604X and +/R604X and R604X/R604X). (**F**) Tricep non-labelled (representing embryonic and Type X) muscle fiber area (+/+=358.8μm^2^, +/R604X=296.2μm^2^, R604X/R604X=267.9μm^2^, *P*<0.0001). (**G**) Percentage of all cells expressing a given muscle fiber type in wild type, *Ighmbp2*^+/R604X^ and *Ighmbp2*^R604X/R604X^ mice (embryonic +/+=6.4, +/R604X=11.5, R604X/R604X=34.6, *P*<0.0001 +/+ and R604X/R604X, *P*=0.0004 +/R604X and R604X/R604X) (Type 2B +/+=49.3, +/R604X=62.3, R604X/R604X=1.6, *P*<0.0001, *P*=0.0117 +/+ and +/R604X) (non-labelled +/+=36.5, +/R604X=27.9, R604X/R604X=85.5, *P*<0.0001). Five mice from each genotype were analyzed. Statistical analyses: one-way ANOVA with Dunnett’s multiple comparisons. Values are expressed as mean.

### Electrophysiology of *Ighmbp2*^R604X/R604X^ mice was consistent with pathology of sciatic nerve, NMJ denervation and hindlimb muscles

There was significant pathology associated with the sciatic nerve, TA and gastrocnemius NMJ denervation, and atrophy of the hindlimb muscles in *Ighmbp2*^R604X/R604X^ mice; therefore, to determine whether functional deficits were present, six electrophysiological parameters were quantitatively measured with relative stimulation of the sciatic nerve and response of the gastrocnemius muscle (Figure 13). The six electrophysiological parameters were: distal latency, negative area, P-P CMAP (peak to peak compound muscle action potential) amplitude, O-P CMAP (onset to peak compound muscle action potential) amplitude, MUNE (motor unit number estimation) and RNS (repetitive nerve stimulation) (Figure 13). Distal latency measures the time it takes for the electrical signal to travel from the stimulation point to the recording point and is reflected by the axon caliber. Negative area is a measurement of the number of muscle fibers activated. P-P CMAP amplitude measures the output from the muscle following maximal stimulation of the nerve and is the difference in voltage between the highest and lowest points (peaks) of the signal. O-P CMAP amplitude measures the strength of the initial nerve to muscle response. MUNE is a calculated estimate of the number of motor units and RNS is how much the CMAP amplitude changes during a set of electrical stimulations. For all six parameters there were significant differences between wild type and *Ighmbp2*^R604X/R604X^ mice (Figure 13A-I). There were no statistical differences between wild type and *Ighmbp2*^+/R604X^ mice (Figure 13). The prolonged distal latency in *Ighmbp2*^R604X/R604X^ mice was consistent with the significant pathology observed in the sciatic nerve including reduced area, myelin thickness and number of myelinated axon fibers. The reduced negative area supports the disease pathology observed in the nerve and hindlimb muscles of *Ighmbp2*^R604X/R604X^ mice. The significantly reduced P-P and O-P CMAP in *Ighmbp2*^R604X/R604X^ mice support the pathology associated with sciatic nerve and NMJ denervation while the reduced MUNE was consistent with the significant NMJ denervation of the hindlimb muscles. Finally, the increased RNS detected in *Ighmbp2*^R604X/R604X^ mice (+/+=4.6% reduction in repetitive stimulation, R604X/R604X=16.5% reduction in repetitive stimulation) was consistent with impaired neuromuscular transmission.

**Figure 13.**
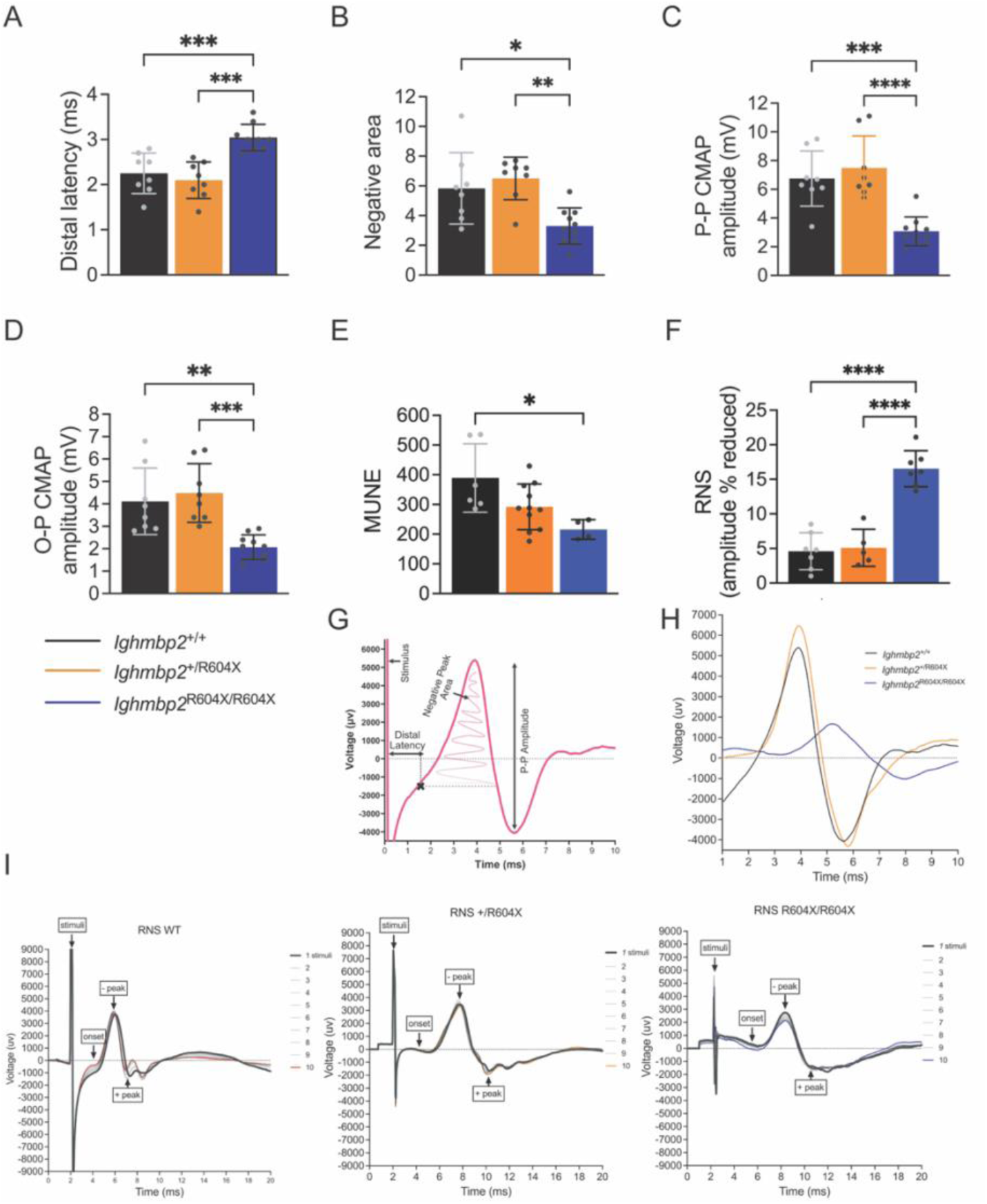
Prolonged distal latency and reduced CMAP in *Ighmbp2*^R604X/R604X^ mice were consistent with pathology of sciatic nerve, NMJ denervation and hindlimb muscles. Wild type (black), *Ighmbp2*^+/R604X^ (orange), and *Ighmbp2*^R604X/R604X^ (blue). (**A**) Distal latency (+/+=2.25 milliseconds, +/R604X=2.10 milliseconds, R604X/R604X=3.04 milliseconds, *P*=0.0001 +/+ and R604X/R604X, *P*=0.0009 +/R604X and R604X/R604X). (**B**) Negative area (+/+=5.84, +/R604X=6.50, R604X/R604X=3.30, *P*=0.0132 +/+ and R604X/R604X, *P*=0.0019 +/R604X and R604X/R604X). (**C**) P-P CMAP amplitude (+/+=6.75 millivolts, +/R604X=7.49 millivolts, R604X/R604X=3.07 millivolts, *P*=0.0005 +/+ and R604X/R604X, *P*<0.0001 +/R604X and R604X/R604X). (**D**) O-P CMAP amplitude (+/+=4.11 millivolts, +/R604X=4.48 millivolts, R604X/R604X=2.07 millivolts, *P*=0.0027 +/+ and R604X/R604X, *P*=0.0005 +/R604X and R604X/R604X). (**E**) MUNE P-P CMAP amplitude (+/+=389 millivolts, +/R604X=292 millivolts, R604X/R604X=216 millivolts, *P*=0.0140 +/+ and R604X/R604X). (**F**) RNS expressed as amplitude percent reduced (+/+=4.60, +/R604X=5.10, R604X/R604X=16.53, *P*<0.0001). (**G**) action potential diagram depicting electrophysiological measurements. (**H**) action potential trace for wild type, *Ighmbp2*^+/R604X^ and *Ighmbp2*^R604X/R604X^ mice. (**I**) RNS trace for wild type, *Ighmbp2*^+/R604X^ and *Ighmbp2*^R604X/R604X^ mice. Statistical analyses: one-way ANOVA with Tukey’s multiple comparisons. Eight to nine mice from each genotype were analyzed. Values are expressed as mean. ms=milliseconds, mV=millivolts, P-P=peak to peak, CMAP=compound muscle action potential, O-P=onset to negative peak, MUNE=motor unit number estimation, RNS=repetitive nerve stimulus.

### ssAAV9-*IGHMBP*2 gene therapy has minimal impact on survival or fitness before death ensues

The rapid decline in survival and fitness of *Ighmbp2*^R604X/R604X^ mice suggested that we would not achieve robust expression of the ssAAV9-*IGHMBP*2 vector to observe significant therapeutic changes; nonetheless, we measured survival, weight and presence of a milk spot in wild type, *Ighmbp2*^+/R604X^, *Ighmbp2*^R604X/R604X^, *Ighmbp2*^R604X/R604X^ + ssAAV9-*IGHMBP2* and *Ighmbp2*^R604X/R604X^ + ssAAV9-*IGHMBP2* + scAAV9-*Abt1* mice. Mean survival was increased three days in *Ighmbp2*^R604X/R604X^ + ssAAV9-*IGHMBP2* mice and one day in *Ighmbp2*^R604X/R604X^ + ssAAV9-*IGHMBP2* + scAAV9-*Abt1* mice over *Ighmbp2*^R604X/R604X^ uninjected controls (Supplemental Figure 5A). Weight did not see significant changes between uninjected and injected *Ighmbp2*^R604X/R604X^ mice; however, *Ighmbp2*^R604X/R604X^ + ssAAV9-*IGHMBP2* mice maintained weight for several days longer prior to weight loss (Supplemental Figure 5B). When the milk spot was scored there was not a significant difference between uninjected and injected *Ighmbp2*^R604X/R604X^ mice; however, consistent with weight, the presence of a milk spot was maintained longer in *Ighmbp2*^R604X/R604X^ + ssAAV9-*IGHMBP2* mice than uninjected controls (Supplemental Figure 5C). In none of these experiments was expression of scAAV9-*Abt1* able to overcome disease within such a short window of time (Supplemental Figure 5A-C). When the presence of the ssAAV9-*IGHMBP2* vector was analyzed by PCR in the lumbar region of the spinal cord, ssAAV9-*IGHMBP2* vector was detected only in spinal cord tissue injected with the vector (Supplemental Figure 5D). Expression of the ssAAV9-*IGHMBP2* vector was measured by qPCR showing that expression of the ssAAV9-*IGHMBP2* vector was present (Supplemental Figure 5E). Importantly, these experiments shed insight into how different *Ighmbp2* mutations impact disease severity and the ability of gene therapy to modify disease.

## Materials and methods

### Generation of the FVB-*Ighmbp2* mice

Animal procedures were carried out in accordance with procedures approved by the NIH and MU Animal Care and Use Committee. Animals were housed using standard animal husbandry with free access to water and food. For all experiments, litter sizes were culled (5-7 pups/litter) to foster maternal care. Due to the short lifespan of *Ighmbp2*^R604X/R604X^ mice, mice were used in all experiments without sex bias. The mouse *Ighmbp2*-R604X mutation corresponds to the human *IGHMBP2*-R605X mutation. The *Ighmbp2*^+/R604X^ mouse was generated in collaboration with the MU Animal Modeling Core using CRISPR technologies. An enhanced-specificity Cas9 (eSPCas9) protein was used to reduce off-target effects. Any predicted off-target site with less than a 2bp mismatch (including DNA or RNA bulges) or with less than 3bp mismatches if no mismatches were in the 12bp seed region of the sgRNA was PCR amplified and sequenced to ensure no erroneous edits were made. The sgRNA was ordered as chemically modified synthetic sgRNA (Synthego). The repair template was chemically synthesized as a 199bp single stranded DNA oligo (ssODN) (IDT). The ssODN was complementary to the non-target strand and contained symmetrical homology arms. The *Ighmbp2*-R604X single guide RNA (sgRNA) sequence: 5′-ATGTTGCTGTTACCCGTGCT −3′. The repair template sequence (sgRNA sequence disrupted by the desired mutation): 5′-ATCCCTGCCTCTCTCATCTCTGGCTTTTCTTCAGGTGAAGTTGGTTTTCTGGCTGAGGACAGGCGGATTAA TGTTGCTGTTACCCGTGCT**<u>TA</u>G**CGGCACGTGGCAGTCATCTGTGATTCCCACACTGTCAACAACCATGCT

TTTTTGAAGACCTTGGTGGATTATTTCACAGAGCATGGGGAGGTACGCACAGCCTTTGA-3′. The TAG indicated in bold is the premature stop codon with <u>TA</u> modified in the repair template. Founder mice genotypes were confirmed by sequencing and none of the founders contained off-site mutations. Founders were outcrossed to FVB wild type mice (Jackson Laboratory) to establish the colony. Animals were extensively backcrossed prior to studies.

### Genotyping

Genotyping of neonatal pups was performed at P0. Genomic DNA was obtained using the DNA isolation protocol from Jackson Labs. The *Ighmbp2*-R604X mutant allele was determined using FWD 5′-TTCAGGTGAAGTTGGTTTTCTGGC-3′ and REV 5’-GTATTCAAAGGCTGTGCGTACCTCC-3′ primers and GoTaq. PCR conditions were step 1: 94°C denaturing for 3:00 minutes, step 2: 30 cycles of 94°C denaturing for 30 seconds, 60°C annealing for 30 seconds, and 68°C extension for 1:00 minute, step 3: final extension was 68°C for 5 minutes. Amplicons for the R604X assay were digested with *Dde*I (NEB) and separated on a 4% agarose gel to differentiate wild type from mutant alleles. Wild type restriction enzyme digestion products were 161bp, 151bp, and 23bp. Mutant restriction enzyme digestion products were 161bp, 115bp, 36bp, and 20bp.

### Motor function assessments

The hindlimb suspension test measured neonate hindlimb muscle strength from P2-P7 as latency to fall. Neonatal mice were suspended from their hindlimbs from the edge of a 50-ml conical tube with gauze padding at the bottom of the tube. The time from suspension until the fall of the mouse was recorded up to thirty seconds. Each mouse was recorded three times daily with the average of the three scores recorded as the daily average.

### Milk spot

The presence of a milk sac/spot was used as an indication of the mouse pup suckling/feeding. Measured on the left side of the pup, the milk sac is fully visible when milk is present. The presence of a milk spot was scored from P0 to P8 when the milk spot became difficult to access due to the presence of hair on non-mutant mice. The score of 2 indicated the presence of a full milk sac, the score of 1 indicated the presence of a partially full milk sac and the score of 0 indicated no milk was observed in the milk sac.

### Head-out plethysmography

Mice were placed in customized head-out pup chambers equipped with warming beds (Data Sciences International/Harvard Bioscience, Holliston, MA) with a low chamber volume for acquiring low amplitude respiratory signals. Gaps surrounding the neck were sealed using 3M Impregum F base and catalyst. Ventilation was assessed at P2. The mice were acclimated to the chamber while breathing room air (21% O_2_ + 0% CO_2_ + 79% N_2_) for 5 minutes before ventilatory measurements were recorded for baseline conditions for an additional 30 minutes. The mice were then challenged by exposing them to 5 minutes of hypercapnia (7% CO_2_, 21% O_2_, balanced N_2_) and subsequently with hypoxia (10.5% O_2_) + hypercapnia (7% CO_2_, balanced N_2_) for 5 minutes (gas concentrations controlled by a gas mixer; CWe, Inc., Ardmore, PA). A pressure calibration signal, ambient pressures, and chamber pressures were utilized for automated calculation of breath-by-breath respiratory parameters [frequency (f-breaths/min), inspiratory time (Ti-sec), expiratory time (Te-sec), tidal volume (VT-mL), minute ventilation (VE-mL/min), peak inspiratory flow (PIF-mL/sec), and peak expiratory flow (PEF-mL-sec)] at 10-s intervals using FinePointe Software (Data Sciences International/Harvard Bioscience, Holliston, MA). VT, VE, and mean inspiratory flow (VT/Ti-mL/sec) were normalized to body weight (per g). For each treatment, five consecutive time points were selected, averaged, and analyzed. Selection of data points was based on consistency between readings (32). Data were rejected if there was evidence of pressure fluctuations caused by gross body movements.

### Lung and milk aspiration

For lungs, pathology was quantified based off the protocols (33, 34). Briefly, three 20x images of 1 lung section per animal were scored blindly. Thickening of the alveolar wall was scored 0-18 points. Each image/mouse was scored 0-6 with 0=no thickening of alveolar wall (1 cell layer thick), 1-2=mild thickening, 3-4=moderate thickening of alveolar wall, 5=extensive thickening of alveolar wall, 6=complete thickening. Hyaline membrane was scored as acellular deposits devoid of hematoxylin staining in alveolar region but stained with eosin stain and was scored 0-6 points. Each image/mouse was scored 0-2 with 0=no acellular debris, 1=partial build up of acellular debris, 2=significant build up of acellular debris. Observed enhanced injury was scored 0-18 points. Each image/mouse was scored 0-6 with 0=no damage, 1=minimum damage, 2=mild damage, 3=moderate damage, 4=pronounced damage, 5=extensive damage, 6=completely damaged. Atelectasis (complete or partial collapse of distal air spaces) was scored 0-12. Each image/mouse was scored 0-4 with 0=no collapse, 1=slight collapse, 2=50% collapse, 3=nearly full 75% collapse, 4=full collapse of distal air spaces. Milk aspiration was determined by immunohistochemistry of milk proteins using pre-absorbed antibody (Biorbyt orb22391) at 1:500. Three to four 10x images of one lung section were scored blindly with five animals per genotype scored. Alveoli were counted with a plus or minus for detection of milk protein (+/+=708 total alveoli scored with 118+ alveoli/mouse, +R604X=683 total alveoli scored with 102+ alveoli/mouse, R604X/R604X=593 total alveoli scored with 91+ aveoli/mouse).

### Phrenic and sciatic nerve dissection and processing

Phrenic and sciatic nerves were fixed with 8% glutaraldehyde in phosphate buffer and embedded in resin (Poly/Bed^®^ 812; catalog:21844–1; Polysciences Inc.). After fixation, nerves were incubated in 2% osmium tetroxide in phosphate buffer for 45 minutes followed by rinses in ascending ethanol concentrations (50%, 70%, 80%, 95%, and 100%) and propylene oxide. Samples were then incubated in a 1:1 propylene oxide: resin mixture for 1 hour, incubated in resin overnight, placed in a resin mold, and cured at 60°C for 8 hours. Semi-thin sections of 1μm were stained with alkaline toluidine blue, cover-slipped with Permount mounting medium (ThermoFisher Scientific), and visualized by light microscopy at 100x magnification (Leica DM5500 B, Leica Microsystems Inc.). Image quantification was performed in a blind manner using the semi-automated MyelTracer software (35). Myelinated fiber counts were counted using ImageJ (FIJI).

### NMJ immunohistochemistry

Whole mount preparations were post-fixed in 4% PFA following muscle dissection. Anti-neurofilament heavy chain (NF-H) (1:2,000, catalog:AB5539, Millipore) and anti-synaptic vesicle 2 (SV2) (1:200, catalog:YE269, Life Technologies) primary antibodies were used. Donkey anti-chicken Alexa Fluor 488 (1:400, catalog:703–545-155, Jackson ImmunoResearch) and goat anti-rabbit Alexa Fluor 488 (1:200, catalog:111–545-003, Jackson ImmunoResearch) secondary antibodies were used to label the axon and synaptic terminal. Acetylcholine receptors were labeled with Alexa Fluor 594-conjugated α-bungarotoxin (1:200, catalog:B13423, Life Technologies). Three individual images were acquired at 20x magnification using a Leica DM5500 B microscope (Leica Microsystem Inc.) and analyzed blindly. Images were analyzed based on the end plate overlap with the synaptic terminal. Endplates with missing overlapping terminals were considered fully denervated, endplates with partial overlap were considered partially denervated, and endplates with complete overlap were considered fully innervated.

### Diaphragm and skeletal muscle immunohistochemistry

Animals were sacrificed and tissues were dissected. Fresh tissue was flash frozen in liquid nitrogen-chilled 2-methyl-2-butene then embedded in OCT compound (Tissue-Tek© OCT Compound, catalog: 4583, Sakura Finetek USA) and flash frozen again. 10μm sections were co-stained with anti-Laminin primary antibody (1:200; catalog: L9393; Millipore Sigma), myosin heavy chain (MyHC) type 1 (1:10, catalog: BA-D5-s, Iowa Developmental Studies Hybridoma Bank), MyHC type 2A (1:50, catalog: SC-71-s, Iowa Developmental Studies Hybridoma Bank), MyHC type 2B (1:5, catalog: BF-F3-s, Iowa Developmental Studies Hybridoma Bank). Secondary antibody for laminin was Goat anti-Rabbit, Alexa Fluor™ 647 (1:500, catalog: A21244, ThermoFisher Scientific), (MyHC) type 1 Goat anti-Mouse, Alexa Fluor™ 350 (1:500, catalog: A21140, ThermoFisher Scientific), MyHC type 2A Goat anti-Mouse, Alexa Fluor™ 555 (1:500, catalog: A21127, ThermoFisher Scientific), and MyHC type 2B Goat anti-Mouse, Alexa Fluor™ 488 (1:500, catalog: A21042, ThermoFisher Scientific). For the analysis of embryonic muscle fiber type, fresh 10 µm histological sections were co-stained with anti-Laminin primary antibody (1:200; catalog: L9393; Millipore Sigma) and eMyHC (embryonic) (1:5, catalog: F1.652, Iowa Developmental Studies Hybridoma Bank). Secondary antibody for laminin was Goat anti-Rabbit, Alexa Fluor™ 647 (1:500, catalog: A21244, ThermoFisher Scientific) and eMyHC Goat anti-Mouse, Alexa Fluor™ 555 (1:500, catalog: A21127, ThermoFisher Scientific). Muscle fiber area and diameter were analyzed in a blinded manner using SMASH, semi-automatic muscle analysis using segmentation of histology (36).

### Hematoxylin and Eosin staining

Fresh lung tissues were harvested and fixed in 4% PFA for 24 hours at 4°C on a shaker prior to freezing and embedding and cryosectioned at 10μm. Tissue slides were fixed with acetic acid alcohol for 5 minutes, rinsed in tap water twice for 15 seconds and then hematoxylin (catalog: 411165000, ThermoFisher Scientific) stained for 1 minute. Slides were then rinsed in ammonia water for 15 secconds, dehydrated in 95% ethanol twice for 15 seconds and eosin Y (catalog: 152885000, ThermoFisher Scientific) stained for 15 seconds. Slides were then rinsed twice for 15 seconds in 95% ethanol, 100% ethanol, and xylene. Tissues were covered with Permount mounting medium (catalog: SP15–500, ThermoFisher Scientific). Three individual images/mouse at were acquired at 10x magnification using a Leica DM5500 B microscope (Leica Microsystem Inc.) and analyzed blindly.

### Electrophysiology studies

Measurements of the right gastrocnemius muscle were recorded following stimulation of the sciatic nerve (37). Briefly, mice were anesthetized with isoflurane using a Somnoflo vaporizer and mouse pup anesthesia cone (2% for induction (150mL/min) and 1.5% (100mL/min for maintenance) and placed on a warming mat set at 37°C. An active ring electrode was placed over the hindlimb gastrocnemius muscle, and a reference ring electrode was placed over the metatarsals of the right hind paw (Alpine Biomed, Skovlunde, Denmark). Spectra 360 electrode gel was applied to decrease impedance (Parker Laboratories, Fairfield, NJ). A common reference electrode was placed around the tail. Two 28-gauge monopolar needle electrodes (Teca, Oxford Instruments Medical, New York, NY) were placed on each side of the sciatic nerve in the region of the proximal thigh. A portable electrodiagnostic system (Cadwell Sierra Summit, Kennewick, WA) was used to stimulate the sciatic nerve (0.1ms pulse, 1–10mA intensity).

### Generation of ssAAV9-*IGHMBP2* virus and ICV injections

The single-stranded AAV9-*IGHMBP2* viral vector has been previously described (28, 38). Viral particles were generated in HEK293T cells (ATCC® CRL-3216™) using 25-kDa polyethyleneimine and purified using three CsCl density-gradient ultracentrifugation steps followed by dialysis against PBS buffer. Number of viral genomes were determined by qPCR using SYBR green (catalog:1902522, Thermo Fisher Scientific). The animals were intracerebral ventricular (icv) injected at P0 and P1 with ssAAV9-*IGHMBP2* and total of 1.3 × 10^11^ viral genomes. Genomic DNA was extracted from the lumbar region of the spinal cord of the injected mice and controls. The presence of the vector was detected by PCR using the following primers: SV40-Forward 5′-GGATGTTGCCTTTACTTCTAGGCC and *IGHMBP2*-Reverse 5′-GCGTCTCTCTCGAGCTCCAG with a melting temperature of 60°C. For qPCR, lumbar spinal cords were dissected from P6 *Ighmbp2*^R604X/R604X^ + ssAAV9*-IGHMBP2*, uninjected *Ighmbp2*^R604X/R604X^ mice, and *Ighmbp2*^+/+^ mice, flash frozen in liquid nitrogen, and stored at –80 °C until processing. Total RNA was isolated using the RNeasy Mini Kit (Qiagen, Cat. No. 74104) according to the manufacturer’s protocol, including on-column DNase I digestion (Qiagen RNase-Free DNase Set, Cat. No. 79254) to eliminate genomic DNA contamination. RNA concentration and purity were determined using a NanoDrop spectrophotometer (Thermo Fisher Scientific). For cDNA synthesis, 2 µg of total RNA was reverse transcribed using SuperScript IV Reverse Transcriptase (Thermo Fisher Scientific, Cat. No. 18090050). Quantitative PCR was performed on a CFX Real-Time PCR Detection System (Bio-Rad) with SYBR Green PCR Master Mix (Applied Biosystems, Cat. No. 4309155). Viral transgene expression was quantified using primers specific for the vector using forward (SV40) 5’-GGATGTTGCCTTTACTTCTAGGCC-3’ and reverse (*IGHMBP2*) 5’-GCGTCTCTCTCGAGCTCCAG-3’primers, with a melting temperature of 60 °C. The quantification was determined by comparison to a standard curve generated from serial dilutions of the target template.

### Statistics

All experiments were performed in at least three biological replicates for reproducibility of data. The statistics analyses performed for each experiment are included within the figure legends using GraphPad Prism. *P* values less than 0.05 were considered statistically significant. The number of animals within cohorts is indicated within the figure legends. Survival analyses were determined using survival data summary. Statistical analyses for weight, nerve analyses, and electrophysiology were determined using one-way ANOVA with Tukey’s multiple comparison analyses. Statistical analyses for muscle fiber type were determined using one-way ANOVA with Dunnett’s multiple comparison analyses. Statistical analyses for milk spot, plethysmography, NMJ, were determined using two-way ANOVA with Tukey’s multiple comparison analyses.

### Study approval

All experimental procedures were approved by the University of Missouri’s Institutional Animal Care and Use Committee and were performed according to the guidelines outlined in the Guide for the Use and Care of Laboratory Animals.

## Data availability

For original raw data, please contact the corresponding author.

## Author contributions

JLT, RM, MW, DPL conducted experiments, acquired data, analyzed data, and edited the manuscript. CS conducted experiments. CLL and NN analyzed data and edited the manuscript. MAL designed research studies, conducted experiments, acquired data, analyzed data, prepared the manuscript, and edited the manuscript. Authorship order of co-first authors was decided by alphabetical order of last names.

## Acknowledgements

We acknowledge the MU Animal Modeling Core, MU Genomics Technology Core, and MU Advanced Light Microscopy Core for assistance with these studies. The Leica ARTOS 3D ultramicrotome used in these studies was funded by NIH S10OD032246. The antibodies used for muscle fiber type characterization were obtained from the Developmental Studies Hybridoma Bank, created by the NICHD of the NIH and maintained at The University of Iowa, Department of Biology, Iowa City, IA 52242. This work was supported by a NIH/NINDS MPI award to C.L. Lorson and M.A. Lorson (1R01NS134816-01). DPL was supported by a Southeastern Conference (SEC) Scholar fellowship. MW was funded by the IMSD/MARC program National Institutes of Health Training grant T34 GM136493.

## Discussion

Disease onset and clinical symptoms vary significantly for patients with the compound heterozygous *IGHMBP2*-R605X mutation from early onset SMARD1 to a milder CMT2S (4, 8, 26). To understand how the *IGHMBP2*-R605X mutation independently impacts disease, we generated the orthologous mutation in mice ( *Ighmbp2*-R604X) and found that *Ighmbp2*^R604X/R604X^ mice present with the most severe symptoms associated with SMARD1, shortened lifespan, failure to thrive, respiratory distress, reduced suckling and motor function deficits. Interestingly, at birth *Ighmbp2*^R604X/R604X^ mice were able to suckle, based on the presence of a milk spot; however, the milk spot quickly diminished suggesting that the ability to suckle became impaired, consistent with SMARD1 disease in humans. Aspiration is a clinical symptom of SMARD1 patients that exacerbates poor respiration and leads to respiratory infections, increased critical care and death. The *Ighmbp2*^R604X/R604X^ mice are the first to demonstrate this functional deficit. Further studies will focus on examining the SMARD1 disease symptoms of suckling and swallowing difficulties to understand how any changes could impact respiration, nutrition and survival. We found that ssAAV9-*IGHMBP2* gene therapy administered at P0 prolonged survival a few days. Our studies demonstrate that the ssAAV9-*IGHMBP2* vector was identified and expressed within the lumbar spinal cord; therefore, we predict robust expression of the ssAAV9-*IGHMBP2* vector was not achieved prior to death. Administration of ssAAV9-*IGHMBP2* at P0 and scAAV9-*Abt1* at P1 did not provide a survival advantage.

While there were no statistical differences between wild type and *Ighmbp2*^+/R604X^ mice in lifespan, weight, presence of a milk spot, plethysmography, electrophysiology, lung pathology and NMJ innervation status of all muscles examined, we did see differences in pathology associated with diaphragm, TA, gastrocnemius and tricep muscle fiber area and phrenic and sciatic axon pathology. It appears that these cellular changes are not sufficiently impactful to result in a physical phenotype, consistent with SMARD1 being a recessive disease.

SMARD1 is a disease associated with severe respiratory distress. P2 *Ighmbp2*^R604X/R604X^ mice demonstrated significant respiratory changes measured from eight parameters. *Ighmbp2*^D564N/D564N^ mice also demonstrated significant respiratory changes with decreased frequency and increased tidal volume and an inability to respond to conditions of hypoxia + hypercapnia suggesting deficits in chemoreception (28). *Ighmbp2*^R604X/R604X^ had decreased frequency and trended towards decreased tidal volume with no response to hypercapnia nor hypoxia + hypercapnia conditions suggesting impairment of chemoreception as well. The lack of a chemoreception response likely leads to an imbalance of blood gases and increased blood pH, adding to the severity of the disease. Abnormal plethysmography and lung pathology supported impairment of air flow in *Ighmbp2*^R604X/R604X^ mice, further contributing to respiratory distress. Changes in the phrenic axons and diaphragm muscle fibers likely contribute significantly to the respiratory changes observed in *Ighmbp2*^R604X/R604X^ mice. Smaller axon fibers with reduced myelin likely reduce the speed of transmission and the efficiency. Much like *Ighmbp2*^D564N/D564N^ and *Ighmbp2*^D564N/H922Y^ mice, there was not substantial NMJ denervation of the diaphragm muscle in *Ighmbp2*^R604X/R604X^ mice suggesting NMJ denervation is not a predictor of respiratory distress in SMARD1 mice (25, 28). While there were no significant changes in the muscle fiber type composition of the diaphragm, all muscle fiber types were significantly smaller, likely contributing towards diaphragm muscle weakness and impaired respiration. These studies suggest that there are multiple pathological changes in *Ighmbp2*^R604X/R604X^ mice that contribute to the respiratory distress and ultimate survival of these SMARD1 mice. Furthermore, our results are consistent with the clinical symptoms associated with severe SMARD1 patients and demonstrate the utility of these mouse models for understanding the underlying cellular changes associated with IGHMBP2 mutations.

To evaluate motor function in these young mice, we performed electrophysiology and examined pathology in the sciatic nerve, NMJ innervation status and muscle pathology of forelimb tricep and hindlimb TA and gastrocnemius muscles. *Ighmbp2*^R604X/R604X^ mice showed significant electrophysiological changes in all parameters analyzed suggesting early gross motor function deficits. *Ighmbp2*^R604X/R604X^ mice electrophysiology was consistent with the reduced sciatic axon myelin thickness and smaller axon fibers impairing transmission time and intensity. Unlike the diaphragm, *Ighmbp2*^R604X/R604X^ forelimb (tricep) muscle showed some NMJ denervation and the hindlimb (TA and gastrocnemius) muscles were severely impacted with NMJ denervation. In contrast to the diaphragm, *Ighmbp2*^R604X/R604X^ forelimb and hindlimb muscle fibers showed significant abnormal pathology and differences in muscle fiber composition. Most notably in *Ighmbp2*^R604X/R604X^ mice, the tricep and TA contained significantly smaller embryonic fibers that represented a significant proportion of the muscle fiber types while Type 2B fibers were significantly underrepresented in comparison to wild type mice. In the tricep and TA, non-labelled fibers (embryonic and Type 2X) constituted over sixty-nine percent of *Ighmbp2*^R604X/R604X^ muscle fibers while only thirty-seven percent in wild type mice. Together the reduced muscle fiber size and changes in muscle fiber composition in *Ighmbp2*^R604X/R604X^ mice are consistent with the electrophysiology data and suggest not only changes in the efficiency of the forelimb and hindlimb muscles but also in the overall function of these muscles.

The *Ighmbp2*^R604X/R604X^ mouse model presents with the most severe SMARD1 clinical symptoms unmasking lung pathology, the loss of the ability to suckle, and aspiration. These studies suggest that *Ighmbp2*^R604X/R604X^ mice likely succumb to death due to severe respiratory distress, and reduced suckle (nutrition), clinical symptoms associated with SMARD1 patients. The IGHMBP2-R605X patient mutation is predicted to generate a truncated IGHMBP2 protein. In a follow-up manuscript, we examine the biochemical properties associated with the IGHMBP2-R605X protein and how the IGHMBP2-R605X mutation alters the association with ABT1. Importantly, we examine how the IGHMBP2-R605X protein interacts with wild type IGHMBP2 protein.

## Notes

Conflict-of-Interest Statement: Lorson <u>Ownership</u> CLL is co-founder and chief scientific officer of Shift Pharmaceuticals. <u>Income</u> CLL has received in excess of $10,000 in income per annum from Shift Pharmaceuticals. <u>Research support</u> Research in the CLL and MAL labs have been supported by sub-awards from Shift Pharmaceuticals (as part of grants from the DOD, CMT Research Foundation, and the NIH). <u>Intellectual property</u> CLL and MU share patents on novel compounds licensed by Shift Pharmaceuticals and planned patents for additional novel compounds. MAL is associated with Shift by family relation.

### Competing Interest Statement

Conflict-of-Interest Statement: Lorson Ownership CLL is co-founder and chief scientific officer of Shift Pharmaceuticals.
Income CLL has received in excess of $10,000 in income per annum from Shift Pharmaceuticals. Research support Research in the CLL and MAL labs have been supported by sub-awards from Shift Pharmaceuticals (as part of grants from the DOD, CMT Research Foundation, and the NIH). Intellectual property CLL and MU share patents on novel compounds licensed by Shift Pharmaceuticals and planned patents for additional novel compounds. MAL is associated with Shift by family relation.

## References

1. Grohmann K, et al. Diaphragmatic spinal muscular atrophy with respiratory distress is heterogeneous, and one form Is linked to chromosome 11q13-q21. Am J Hum Genet. 1999;65(5):1459–1462.

2. Viollet L, et al. Mapping of autosomal recessive chronic distal spinal muscular atrophy to chromosome 11q13. Ann Neurol. 2002;51(5):585–592.

3. Grohmann K, et al. Mutations in the gene encoding immunoglobulin mu-binding protein 2 cause spinal muscular atrophy with respiratory distress type 1. Nat Genet. 2001;29(1):75–77.

4. Guenther UP, et al. Clinical and mutational profile in spinal muscular atrophy with respiratory distress (SMARD): defining novel phenotypes through hierarchical cluster analysis. Hum Mutat. 2007;28(8):808–815.

5. Rudnik-Schoneborn S, et al. Long-term observations of patients with infantile spinal muscular atrophy with respiratory distress type 1 (SMARD1). Neuropediatrics. 2004;35(3):174–182.

6. Porro F, et al. The wide spectrum of clinical phenotypes of spinal muscular atrophy with respiratory distress type 1: a systematic review. J Neurol Sci. 2014;346(1-2):35–42.

7. Kaindl AM, et al. Spinal muscular atrophy with respiratory distress type 1 (SMARD1). J Child Neurol. 2008;23(2):199–204.

8. Cottenie E, et al. Truncating and missense mutations in IGHMBP2 cause Charcot-Marie Tooth disease type 2. Am J Hum Genet. 2014;95(5):590–601.

9. Schottmann G, et al. Recessive truncating IGHMBP2 mutations presenting as axonal sensorimotor neuropathy. Neurology. 2015;84(5):523–531.

10. Wagner JD, et al. Autosomal recessive axonal polyneuropathy in a sibling pair due to a novel homozygous mutation in IGHMBP2. Neuromuscul Disord. 2015;25(10):794–799.

11. Liu L, et al. IGHMBP2-related clinical and genetic features in a cohort of Chinese Charcot-Marie-Tooth disease type 2 patients. Neuromuscul Disord. 2017;27(2):193–199.

12. Perlick HA, et al. Mammalian orthologues of a yeast regulator of nonsense transcript stability. Proc Natl Acad Sci U S A. 1996;93(20):10928–10932.

13. Bohnsack KE, et al. Cellular functions of eukaryotic RNA helicases and their links to human diseases. Nat Rev Mol Cell Biol. 2023;24(10):749–769.

14. Chen YZ, et al. DNA/RNA helicase gene mutations in a form of juvenile amyotrophic lateral sclerosis (ALS4). Am J Hum Genet. 2004;74(6):1128–1135.

15. Moreira MC, et al. Senataxin, the ortholog of a yeast RNA helicase, is mutant in ataxia-ocular apraxia 2. Nat Genet. 2004;36(3):225–227.

16. Guenther UP, et al. IGHMBP2 is a ribosome-associated helicase inactive in the neuromuscular disorder distal SMA type 1 (DSMA1). Hum Mol Genet. 2009;18(7):1288–1300.

17. Fairman-Williams ME, et al. SF1 and SF2 helicases: family matters. Curr Opin Struct Biol. 2010;20(3):313–324.

18. Chen NN, et al. Evidence for regulation of transcription and replication of the human neurotropic virus JCV genome by the human S(mu)bp-2 protein in glial cells. Gene. 1997;185(1):55–62.

19. Vadla GP, et al. ABT1 modifies SMARD1 pathology via interactions with IGHMBP2 and stimulation of ATPase and helicase activity. JCI Insight. 2023;8(2).

20. de Planell-Saguer M, et al. Biochemical and genetic evidence for a role of IGHMBP2 in the translational machinery. Hum Mol Genet. 2009;18(12):2115–2126.

21. Fukita Y, et al. The human S mu bp-2, a DNA-binding protein specific to the single-stranded guanine-rich sequence related to the immunoglobulin mu chain switch region. J Biol Chem. 1993;268(23):17463–17470.

22. Park J, et al. IGHMBP2 deletion suppresses translation and activates the integrated stress response. Life Sci Alliance. 2024;7(8).

23. Prusty AB, et al. RNA helicase IGHMBP2 regulates THO complex to ensure cellular mRNA homeostasis. Cell Rep. 2024;43(2):113802.

24. Vadla GP, et al. The contribution and therapeutic implications of IGHMBP2 mutations on IGHMBP2 biochemical activity and ABT1 association. Biochim Biophys Acta Mol Basis Dis. 2024;1870(4):167091.

25. Ricardez Hernandez SM, et al. Ighmbp2 mutations and disease pathology: Defining differences that differentiate SMARD1 and CMT2S. Exp Neurol. 2025;383:115025.

26. Grohmann K, et al. Infantile spinal muscular atrophy with respiratory distress type 1 (SMARD1). Ann Neurol. 2003;54(6):719–724.

27. Trujillano D, et al. Clinical exome sequencing: results from 2819 samples reflecting 1000 families. Eur J Hum Genet. 2017;25(2):176–182.

28. Smith CE, et al. The Ighmbp2D564N mouse model is the first SMARD1 model to demonstrate respiratory defects. Hum Mol Genet. 2022;31(8):1293–1307.

29. Schiaffino S, and Moretti I. Changes in skeletal muscle fiber types induced by chronic kidney disease. Kidney Int. 2015;88(2):412.

30. Schiaffino S, and Reggiani C. Fiber types in mammalian skeletal muscles. Physiol Rev. 2011;91(4):1447–1531.

31. Westerblad H, et al. Skeletal muscle: energy metabolism, fiber types, fatigue and adaptability. Exp Cell Res. 2010;316(18):3093–3099.

32. Lind LA, et al. Intralingual Administration of AAVrh10-miR(SOD1) Improves Respiratory But Not Swallowing Function in a Superoxide Dismutase-1 Mouse Model of Amyotrophic Lateral Sclerosis. Hum Gene Ther. 2020;31(15-16):828–838.

33. Silva IAN, et al. A Semi-quantitative Scoring System for Green Histopathological Evaluation of Large Animal Models of Acute Lung Injury. Bio Protoc. 2022;12(16).

34. Karpinski BA, et al. Dysphagia and disrupted cranial nerve development in a mouse model of DiGeorge (22q11) deletion syndrome. Dis Model Mech. 2014;7(2):245–257.

35. Kaiser T, et al. MyelTracer: A Semi-Automated Software for Myelin g-Ratio Quantification. eNeuro. 2021;8(4).

36. Smith LR, and Barton ER. SMASH - semi-automatic muscle analysis using segmentation of histology: a MATLAB application. Skelet Muscle. 2014;4:21.

37. Arnold WD, et al. Electrophysiological Motor Unit Number Estimation (MUNE) Measuring Compound Muscle Action Potential (CMAP) in Mouse Hindlimb Muscles. J Vis Exp. 2015(103).

38. Shababi M, et al. Rescue of a Mouse Model of Spinal Muscular Atrophy With Respiratory Distress Type 1 by AAV9-IGHMBP2 Is Dose Dependent. Mol Ther. 2016;24(5):855–866.

