## Supplemental Figures for "The *Ighmbp2*-R604X mouse recapitulates the severe SMARD1 clinical symptoms of aspiration, respiratory and feeding deficits"

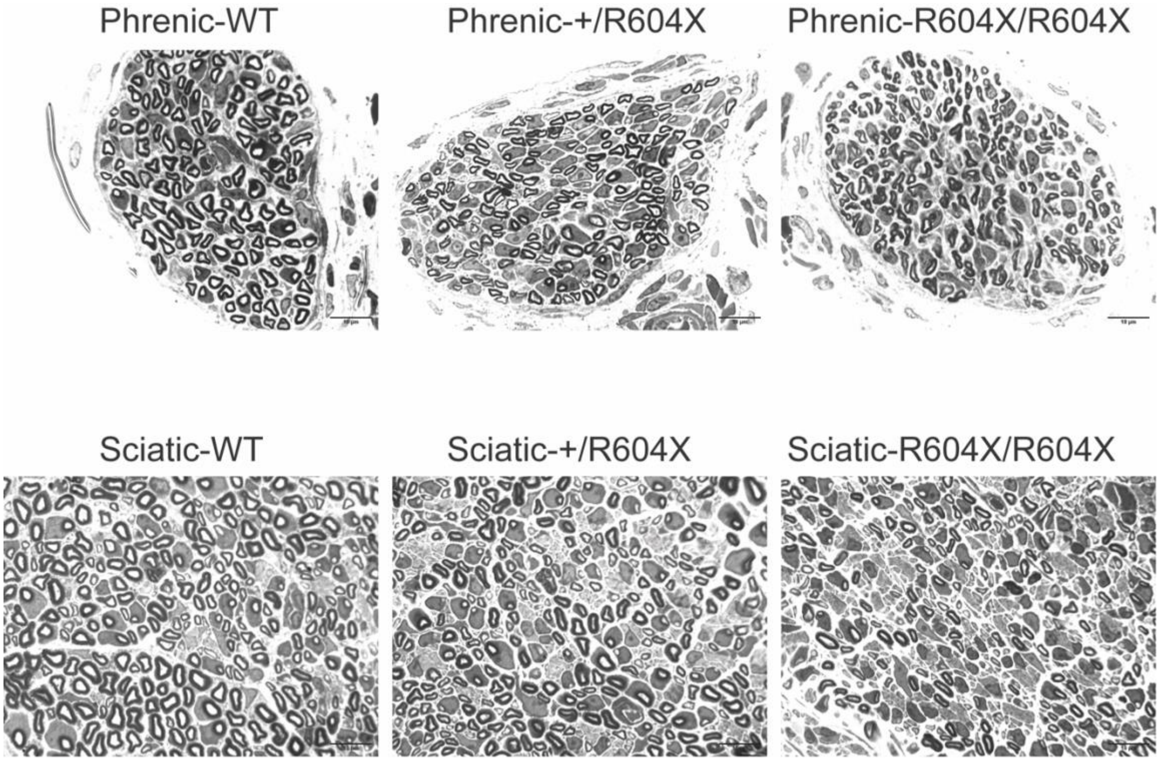


**Supplemental Figure 1. Representative images of phrenic and sciatic nerves.** Representative images of axons of the phrenic and sciatic nerves of wild type (WT), *Ighmbp2*$^{+/R604X}$ and *Ighmbp2*$^{R604X/R604X}$ mice.

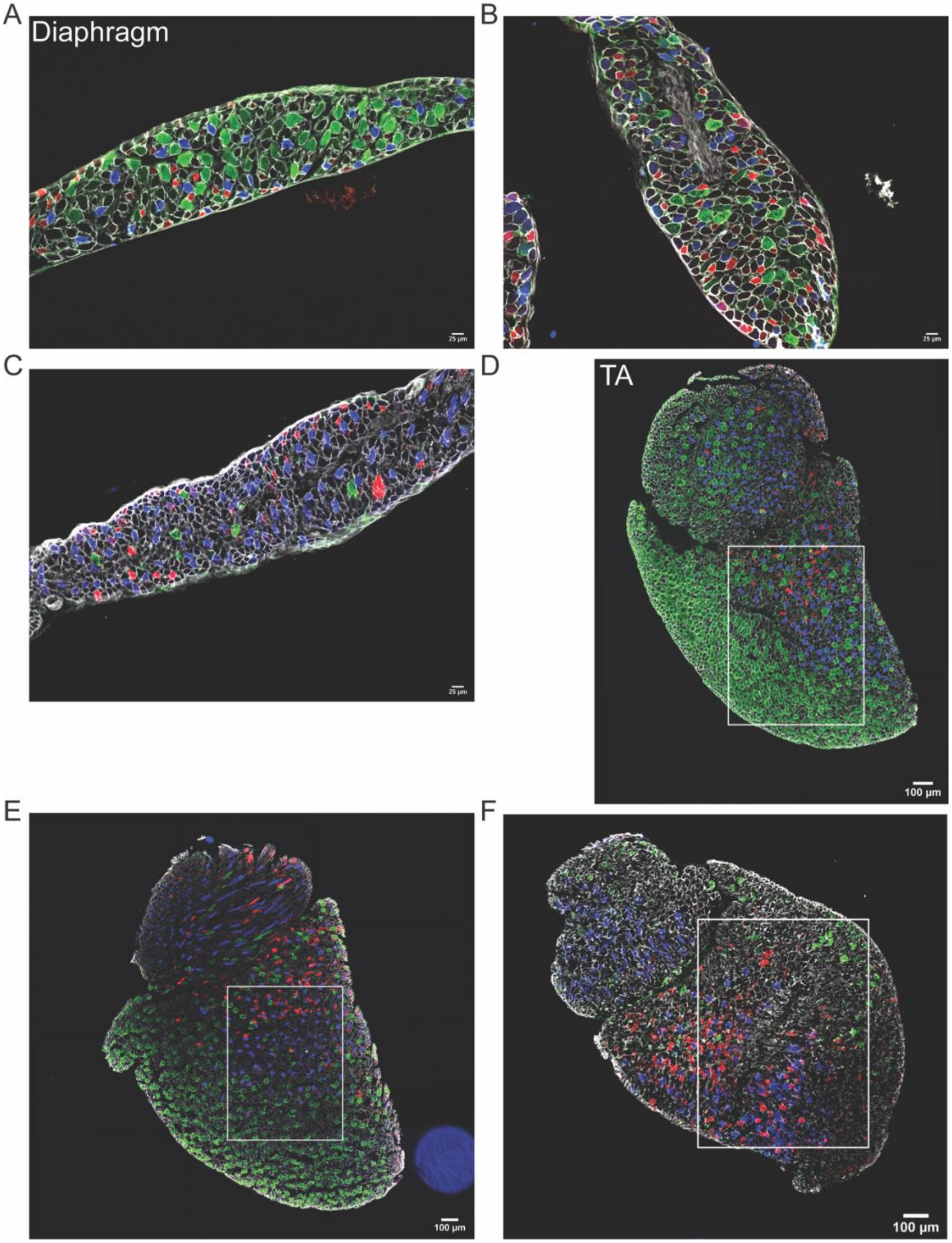


**Supplemental Figure 2. Representative images of the diaphragm and tibialis anterior (TA) muscle fiber types.** Representative images of muscle fibers of the diaphragm and TA of wild type, *Ighmbp2*$^{+/R604X}$ and *Ighmbp2*$^{R604X/R604X}$ mice. (A) Wild type diaphragm muscle. (B) *Ighmbp2*$^{+/R604X}$ diaphragm muscle. (C) *Ighmbp2*$^{R604X/R604X}$ diaphragm muscle. (D) Wild type tibialis anterior (TA) muscle. (E) *Ighmbp2*$^{+/R604X}$ TA muscle. (F) *Ighmbp2*$^{R604X/R604X}$ TA muscle. The square indicates the area that was scored. Blue represents slow twitch type 1 fibers, red represents fast twitch Type 2A fibers, green represents fast twitch Type 2B and no coloring represents embryonic or fast twitch Type 2X. Each muscle fiber was identified using anti-laminin antibodies.

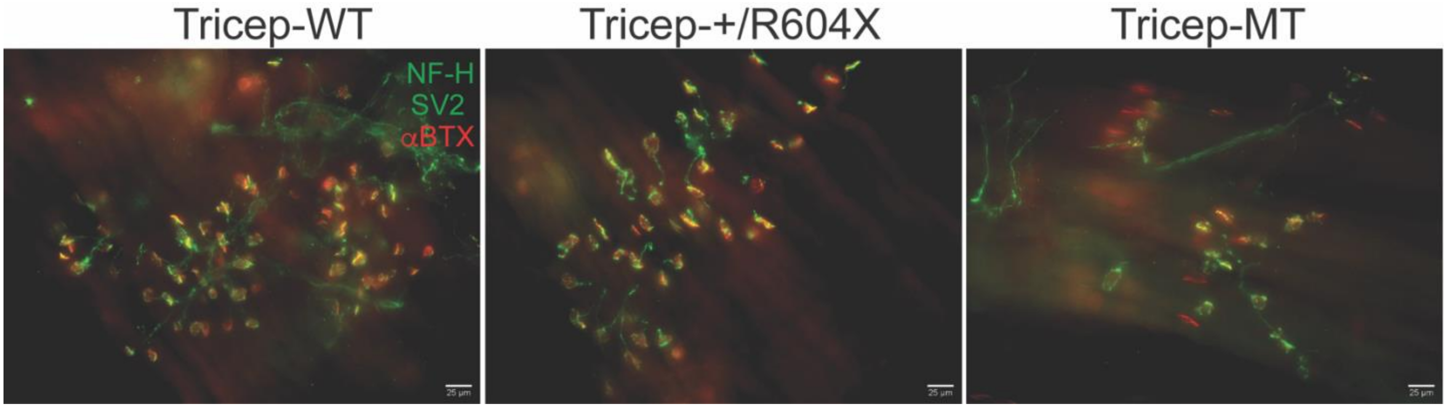


**Supplemental Figure 3: Representative images of NMJ of the tricep muscle.** Representative images of neuromuscular junctions (NMJ) of the tricep of wild type, *Ighmbp2*$^{+/R604X}$ and *Ighmbp2*$^{R604X/R604X}$ mice. Green represents neurofilament heavy (NF-H) and synaptic vesicle 2 (SV2) immunostaining while red represents bungarotoxin 594 immunofluorescence (αBTX). WT=wild type mice, MT=*Ighmbp2*$^{R604X/R604X}$ mice.

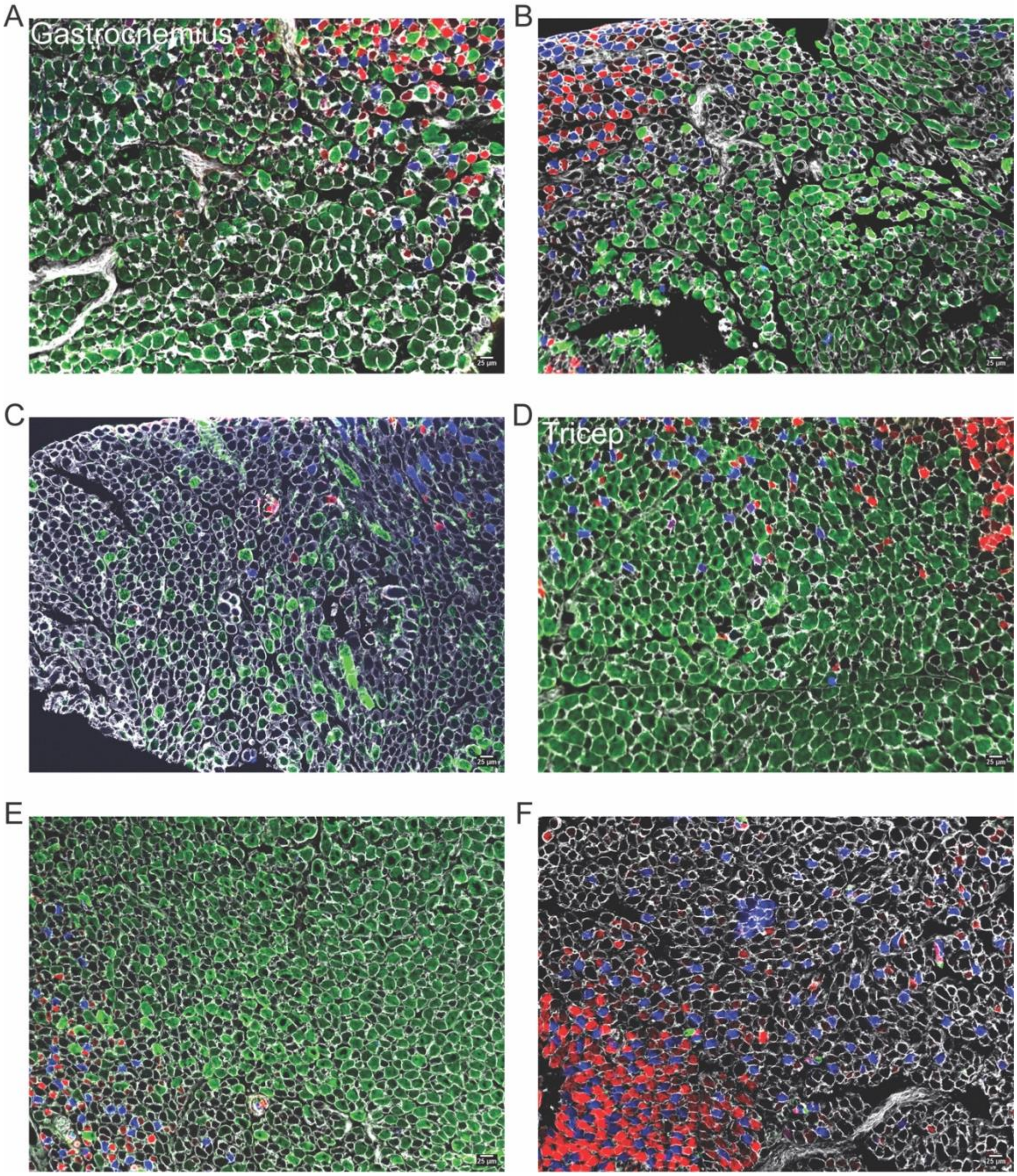


**Supplemental Figure 4. Representative images of the gastrocnemius and tricep muscle fiber types.** Representative images of muscle fibers of the gastrocnemius and tricep of wild type, *Ighmbp2*$^{+/R604X}$ and *Ighmbp2*$^{R604X/R604X}$ mice. (A) Wild type gastrocnemius. (B) *Ighmbp2*$^{+/R604X}$ gastrocnemius. (C) *Ighmbp2*$^{R604X/R604X}$ gastrocnemius. (D) Wild type tricep. (E) *Ighmbp2*$^{+/R604X}$ tricep. (F) *Ighmbp2*$^{R604X/R604X}$ tricep. Blue represents slow twitch Type 1 fibers, red represents fast twitch Type 2A fibers, green represents fast twitch Type 2B and no coloring represents embryonic or fast twitch Type 2X. Each muscle fiber was identified using anti-laminin antibodies.

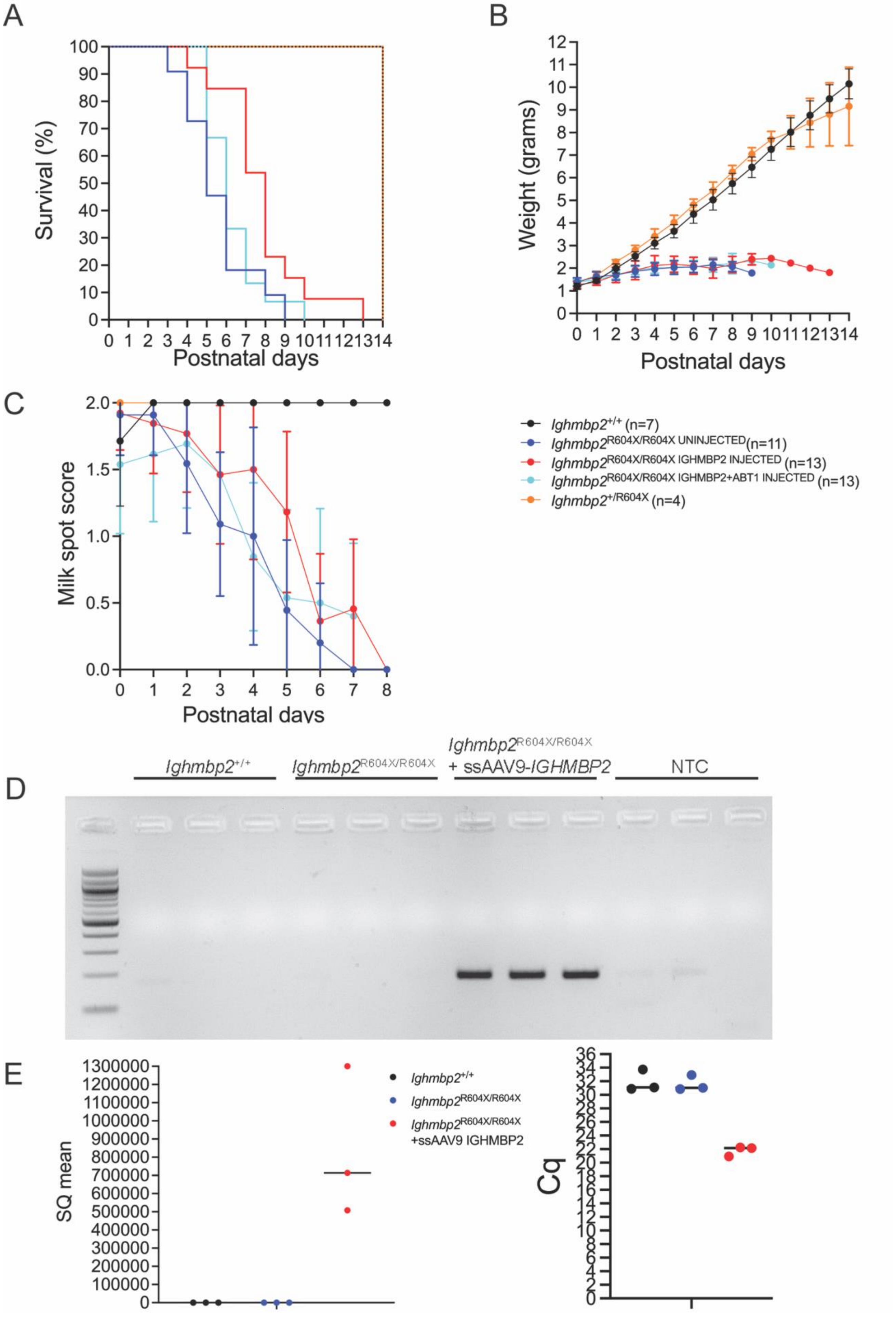
A
Survival (%)
Postnatal days
B
Weight (grams)
Postnatal days
C
Milk spot score
Postnatal days
Ighmbp2+/+ (n=7)
Ighmbp2R604X/R604X UNINJECTED(n=11)
Ighmbp2R604X/R604X IGHMBP2 INJECTED (n=13)
Ighmbp2R604X/R604X IGHMBP2+ABT1 INJECTED (n=13)
Ighmbp2+/R604X (n=4)
D
Ighmbp2+/+
Ighmbp2R604X/R604X
Ighmbp2R604X/R604X + ssAAV9-IGHMBP2
NTC
E
SQ mean
Ighmbp2+/+
Ighmbp2R604X/R604X
Ighmbp2R604X/R604X +ssAAV9 IGHMBP2
Cq

**Supplemental Figure 5. ssAAV9-*IGHMBP*2 gene therapy has slight impact on survival and fitness before death ensues.** Wild type (black), *Ighmbp2*$^{+/R604X}$ (orange), and *Ighmbp2*$^{R604X/R604X}$ (blue), *Ighmbp2*$^{R604X/R604X}$ + ssAAV9-*IGHMBP2* (red), and *Ighmbp2*$^{R604X/R604X}$ + ssAAV9-*IGHMBP2* + scAAV9-*Abt-1* (turquoise). A study of survival, weight and milk spot on same cohorts for fourteen days. (**A**) Survival curve for wild type, *Ighmbp2*$^{+/R604X}$, *Ighmbp2*$^{R604X/R604X}$, *Ighmbp2*$^{R604X/R604X}$ + ssAAV9-*IGHMBP2*, and *Ighmbp2*$^{R604X/R604X}$ + ssAAV9-*IGHMBP2* + scAAV9-*Abt-1* mice followed for fourteen days. Mean survival was fourteen days for wild type and *Ighmbp2*$^{+/R604X}$ mice while the mean survival for *Ighmbp2*$^{R604X/R604X}$ mice was five days ($P$<0.0001). Mean survival for *Ighmbp2*$^{R604X/R604X}$ + ssAAV9-*IGHMBP2* mice was eight days and *Ighmbp2*$^{R604X/R604X}$ + ssAAV9-*IGHMBP2* + scAAV9-*Abt-1* mice six days. (**B**) Weight measured for fourteen days. The mean weight was wild type=5.28 grams, *Ighmbp2*$^{+/R604X}$=5.41 grams, *Ighmbp2*$^{R604X/R604X}$=1.87 grams, *Ighmbp2*$^{R604X/R604X}$ + ssAAV9-*IGHMBP2*=1.98 grams, and *Ighmbp2*$^{R604X/R604X}$ + ssAAV9-*IGHMBP2* + scAAV9-*Abt-1*=1.94 grams. (**C**) Presence of milk spot followed to P8. Score of 2=full milk spot, 1=partial milk spot, 0=no milk spot present. Mean wild type=2.0, *Ighmbp2*$^{+/R604X}$=2.0, *Ighmbp2*$^{R604X/R604X}$=0.9, *Ighmbp2*$^{R604X/R604X}$ + ssAAV9-*IGHMBP2*=1.2, and *Ighmbp2*$^{R604X/R604X}$ + ssAAV9-*IGHMBP2* + scAAV9-*Abt-1*=1.1. There was not a statistical difference in survival, weight or milk spot between *Ighmbp2*$^{R604X/R604X}$, *Ighmbp2*$^{R604X/R604X}$ + ssAAV9-*IGHMBP2*, or *Ighmbp2*$^{R604X/R604X}$ + ssAAV9-*IGHMBP2* + scAAV9-*Abt-1* mice. **(D)** PCR analyses of wild type, *Ighmbp2*$^{R604X/R604X}$ uninjected and *Ighmbp2*$^{R604X/R604X}$ + ssAAV9-*IGHMBP2* injected mice. DNA was extracted from the lumbar spinal cord. Lane 1= NEB 1kb DNA marker. Primers used were to the viral vector and human *IGHMBP2* cDNA. **(E)** qPCR analyses of wild type, *Ighmbp2*$^{R604X/R604X}$ uninjected and *Ighmbp2*$^{R604X/R604X}$ + ssAAV9-*IGHMBP2* injected mice. RNA was extracted from the lumbar spinal cord. Primers used were to the viral vector and human *IGHMBP2* cDNA. Statistical analyses: Survival data summary for survival, one-way ANOVA with Tukey's multiple comparison for weight. Values are expressed as mean, n=number of mice, NTC=no template control.
